# A Curated Compendium of Transcriptomic Data for the Exploration of Neocortical Development

**DOI:** 10.1101/2024.02.26.581612

**Authors:** Shreyash Sonthalia, Ricky S. Adkins, Joshua Orvis, Guangyan Li, Xoel Mato Blanco, Alex Casella, Jinrui Liu, Genevieve Stein-O’Brien, Brian Caffo, Ronna Hertzano, Anup Mahurkar, Jesse Gillis, Jonathan Werner, Shaojie Ma, Nicola Micali, Nenad Sestan, Pasko Rakic, Gabriel Santpere, Seth A. Ament, Carlo Colantuoni

## Abstract

Vast quantities of multi-omic data have been produced to characterize the development and diversity of cell types in the cerebral cortex of humans and other mammals. To more fully harness the collective discovery potential of these data, we have assembled gene-level transcriptomic data from 188 published studies of neocortical development, including the transcriptomes of ∼30 million single-cells, extensive spatial transcriptomic experiments and RNA sequencing of sorted cells and bulk tissues: nemoanalytics.org/landing/neocortex. Applying joint matrix decomposition (SJD) to mouse, macaque and human data in this collection, we defined transcriptome dynamics that are conserved across mammalian neurogenesis and which elucidate the evolution of outer, or basal, radial glial cells. Decomposition of adult human neocortical data identified layer-specific signatures in mature neurons and, in combination with transfer learning methods in NeMO Analytics, enabled the charting of their early developmental emergence and protracted maturation across years of postnatal life. Interrogation of data from cerebral organoids demonstrated that while broad molecular elements of *in vivo* development are recapitulated *in vitro*, many layer-specific transcriptomic programs in neuronal maturation are absent. We invite computational biologists and cell biologists without coding expertise to use NeMO Analytics in their research and to fuel it with emerging data (carlocolantuoni.org).

## Main

Expansion of the neocortex has been dramatic in primates, most recently elevating cognitive abilities in the human lineage to extraordinary heights (PMID: 19763105). The neurons of the cortex are born and assembled into a conserved laminar architecture during prenatal development (PMID: 3291116; PMID: 26796689; PMID: 29170230). Single cell transcriptomic and epigenomic analyses have been used to generate atlases of mammalian neocortical development (PMID: 30545854; PMID: 34616070; PMID: 34390642; PMID: 36509746; PMID: 37758766; PMID: 34163074; PMID: 34321664; PMID: 37824652) and cerebral organoid models (PMID: 30735633; PMID: 30545853; PMID: 36179669). Two central challenges limit the current discovery potential of these data in the pursuit of a deeper understanding of neocortical development.

The first challenge is that individual datasets cover only a portion of regions, times and species of interest, and are resident in unlinked databases with incomplete metadata, necessitating laborious collection, processing and detailed metadata curation to be interrogated together. Raw and processed data are generally deposited in publicly accessible repositories, but there are limited tools to find, access and assemble data across repositories. Current efforts to bring these data together enable visualization of data from a large number of studies, but only via the focused exploration of individual datasets one-at-a-time (e.g. CellxGene: cellxgene.cziscience.com; Single Cell Portal: singlecell.broadinstitute.org; UCSC Cell Browser: cells.ucsc.edu). To compliment these resources, we have assembled a comprehensive, curated compendium of bulk, microdissected, sorted, spatial, and single-cell transcriptomic data spanning pre- and postnatal ages in mouse, primate and human, as well as *in vitro* stem cell models at nemoanalytics.org/landing/neocortex (Table 1) for the simultaneous analysis of many neocortical datasets.

**Table 1:**
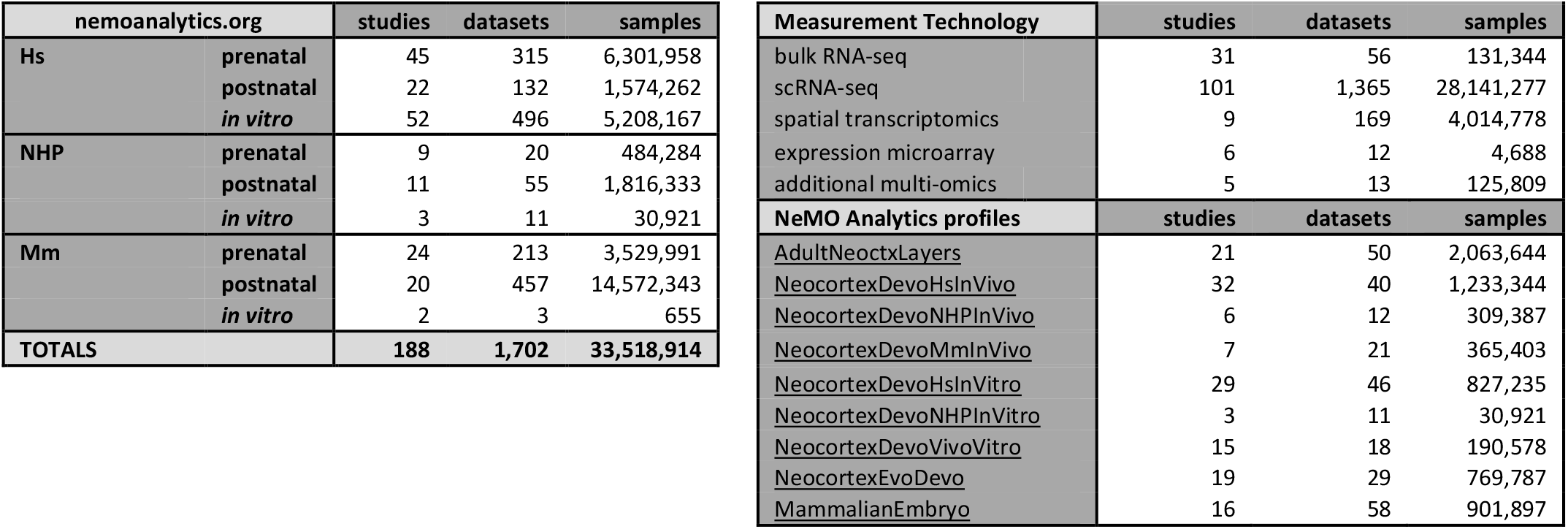
Summary of data resources in nemoanalytics.org. Table contains counts of studies (unique Pubmed IDs), datasets (data matrices) and samples (bulk samples or individual cells) across species and *in vivo* (pre- and post-natal) or *in vitro* studies (full list [HERE](https://docs.google.com/spreadsheets/d/1xag_73Y1xydcCUnDYwn8rlF2rEbbMdTw/)). “Measurement Technologies” section contains counts for different data modalities. “NeMO Analytics profiles” section contains links to collections of highlighted classes of studies and a summary of resources in each. The “MammalianEmbryo” profile contains data from time points before the brain has formed as a defined organ, with datasets relevant to the emergence of the neural lineage and telencephalic progenitors from the pluripotent epiblast.

The second challenge is that standard data analysis pipelines are insufficient to identify common molecular mechanisms across large numbers of diverse datasets. Multi-omic data integration methods have been widely successful in combining datasets containing uniform data modalities and cell types (PMID: 34062119, PMID: 31178122, PMID: 31740819). Here, we apply structured joint decomposition (SJD; doi.org/10.1101/2022.11.07.515489) to selected subsets of the NeMO Analytics data collection to define robust shared dynamics across heterogeneous, but biologically linked gene expression experiments. We uncover both conserved mammalian and primate-specific transcriptome dynamics in neurogenesis and neuronal maturation. Further, we apply transfer learning approaches implemented in NeMO analytics (PMID: 32167521, doi.org/10.1101/2022.11.07.515489) to explore these transcriptome dynamics across the compendium of neocortical data, enabling broad ranging molecular perspectives across evolution and developmental time and space. These resources can be freely utilized by the research community (with and without coding expertise) to explore additional aspects of cortical development, and our approach is readily extensible to other areas of biomedical research.

### NeMO Analytics: A comprehensive transcriptomic data exploration environment for neocortical development

The Neuroscience Multi-Omic (NeMO) Analytics platform (nemoanalytics.org) is designed for cell biologist-friendly visualization and analysis of many transcriptomics datasets in parallel. In NeMO Analytics, we assembled a comprehensive collection of gene-tabulated transcriptomic and additional multi-omic datasets focused on excitatory neocortical neurogenesis and neuronal maturation (Table 1, Figure 1A). The compendium incorporates data from 188 published studies, including the transcriptomes of ∼30 million single-cells, >4 million spatial transcriptomic positions, and >130,000 samples from RNA sequencing of sorted cells and bulk tissues (full list of datasets [HERE](https://docs.google.com/spreadsheets/d/1xag_73Y1xydcCUnDYwn8rlF2rEbbMdTw/)). Researchers can explore measurements of 1] individual genes, e.g. the expression of the EOMES gene, marking neurogenic intermediate progenitor cells across a collection of *in vivo* RNA-seq experiments in human neocortical development (NeMOlink01), or 2] gene signatures, e.g. the summed expression of genes associated with the S or G2M phases of the cell cycle, to distinguish subsets of cycling progenitors across studies in cerebral organoids (NeMOlink02). In addition to adding their own novel datasets to NeMO Analytics, researchers can upload simple gene lists of interest or more complex gene signatures with weights (e.g. from PCA or NMF) to explore specific transcriptomic dimensions across the data collections in NeMO Analytics (Figure 1A and Methods).

**Figure 1:**
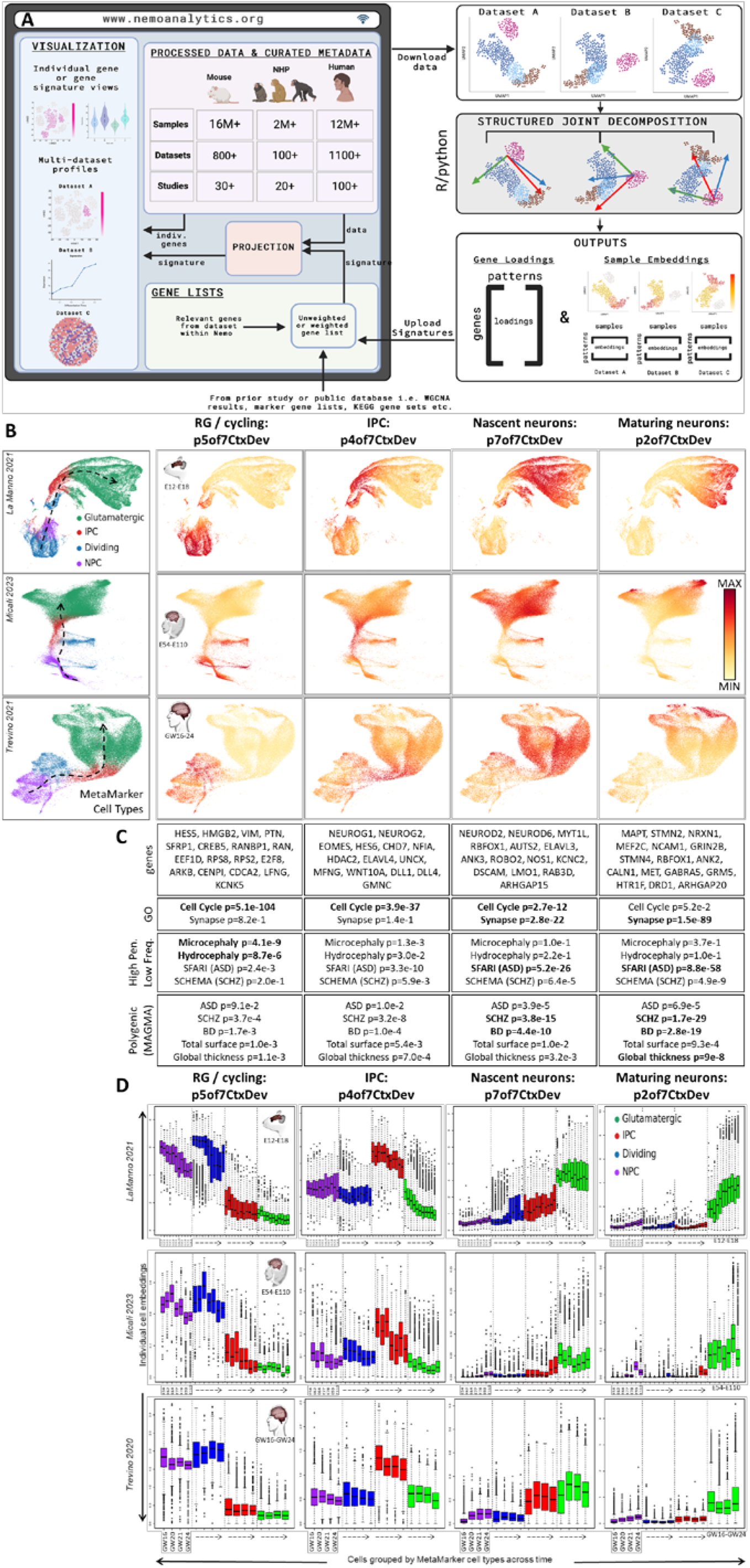
Joint decomposition of scRNA-seq data in mouse, macaque and human neocortical neurogenesis. **A]** Schematic of NeMO Analytics data resources employed in conjunction with joint decomposition and transfer learning approaches. This is an outline of specific analyses in this report as well as a description of a general approach we invite others to take on. Analysis-ready datasets and detailed sample metadata can be downloaded from NeMO Analytics and analyzed offline. Elements learned from joint decomposition can then be uploaded to NeMO Analytics to explore their dynamics across the broad data collection. In this flow, the offline analysis could be any exploratory technique applied to mutli-omics data matrices that produces gene signatures in the form of simple lists or quantitative loadings, e.g. PCA, clustering, or differential expression analysis. **B]** Assembled screen captures from the online NeMO Analytics multi-omic data exploration environment, displaying UMAP representations of scRNA-seq datasets spanning mid-gestational excitatory neocortical neurogenesis in mouse (PMID: 34321664), macaque (PMID: 37824652) and human (PMID: 34390642) development colored by consensus MetaMarker analysis (legend in left-most column). Single-cell embeddings for each of 4 jointNMF patterns selected from a set of 7 (**p7CtxDev**, Figure S1 and NeMOlink03), displayed as color gradients across UMAP plots (columns to right of cell type legend). The jointNMF decomposition produces a single gene loading matrix which underlies these 3 sets of sample embeddings. These gene loadings are used in enrichment analyses in panel C to explore these patterns. Dashed arrows indicate approximate neurogenic trajectory in each species. Individual genes can also be explored in these datasets at NeMOlink04. **C]** Selected genes from the top 0.2% of the gene loadings for each pattern (Table S1 contains all loadings). Genes with highest loadings in p7of7CtxDev include pro-neural genes, while genes encoding proteins that make up physical elements of a neuron are highest in p2of7CtxDev, suggesting that p7 is a transcriptomic program dedicated to becoming a neuron, while p2 represents the neuronal state itself. Selected Gene Ontology (GO) and genetic enrichments in each pattern’s gene loadings are also listed (**BOLD** indicates where hits of greatest significance occur for the different phenotypes). Table S1 contains the full list of enrichments across all 7 patterns. ASD=autism spectrum disorder, SCHZ=schizophrenia, BD=bipolar disorder. **D]** Boxplots of cell embeddings from each of the 4 patterns separated by species, across time and further by MetaMarker-defined cell type labels.

### Transcriptomic dissection of mid-gestation mammalian neocortical neurogenesis via joint matrix decomposition

To leverage a focused subset of this data collection employing joint decomposition approaches, we assembled scRNA-seq data spanning the excitatory neurogenic trajectory in mid-gestational neocortical development in mouse (PMID: 34321664), macaque (PMID: 37824652) and human (PMID: 34390642). In order to first establish a coarse consensus cell labeling across mammalian neocortical development, we used composite expression of “MetaMarkers” (PMID: 37034757), cell type markers that are robust across many studies spanning cortical regions and developmental time (Figure 1B, colored legend, Figure S1A&B, and Methods). Independent of these cell type calls, we applied the jointNMF matrix decomposition algorithm from our SJD package (doi.org/10.1101/2022.11.07.515489) to define shared dimensions of variation resident within all the three input matrices (Methods). Figure 1B depicts 4 of 7 shared transcriptomic patterns that were defined by this approach (patterns from this decomposition will be referred to using the “p7CtxDevo” suffix). These patterns define conserved transcriptomic phases of indirect neocortical neurogenesis spanning distinct progenitor and neuronal states. Patterns p5 and p4of7CtxDev correspond to radial glia cell (RGC) and intermediate progenitor cell (IPC) enriched patterns, respectively. The apparent sequential arrangement of neuronal patterns p7 and p2 in all three species, along with genes highly weighted in these patterns (Figure 1C and Table S1) suggests that p7 is a transient proneural program expressed in nascent neurons, while p2 reflects further neuronal maturation.

We interrogated these conserved transcriptomic elements of neocortical neurogenesis for association with polygenic risk known to play central roles in complex human brain disease using gene level summaries of recent genome-wide association studies in neuropsychiatric disease and brain structure (Methods). Consistent with many recent observations of their broad polygenic nature (PMID: 31464996; PMID: 37853064; PMID: 31907381), the analysis of genome-wide risk yielded strongest associations for schizophrenia and bipolar disorder with the neuronal transcriptome patterns p7 and p2of7CtxDev (Figure 1C). Interestingly, the later neuronal pattern p2 also showed significant association with cortical structure phenotypes. To explore high penetrance, low frequency genetic variation linked to brain disease in these patterns, we conducted enrichment analysis on gene loadings using lists of genes discovered in genome sequencing studies of disease. This revealed that diseases which disrupt the gross structure of the brain, including microcephaly (PMID: 33077954) and hydrocephaly (PMID: 26022163; PMID: 28951247; PMID: 32038172; PMID: 32103185; PMID: 29799801), are associated with p5 that is high in RGCs and especially in cycling progenitors of the developing cortex, consistent with additional recent observations (PMID: 38915580). Genes harboring high penetrance, low frequency variants linked to neuropsychiatric and neurodevelopmental disorders were strongly enriched in the neuronal patterns p7 and p2, with ASD having particularly strong associations. This is consistent with many observations indicating that low-frequency, high-penetrance *de novo* variants play a key role in this disorder (PMID: 32668441; PMID: 35440779). These findings indicate that the distinct genetic architectures underlying these different cortical disorders play out in particular elements of the neurogenic transcriptome (see Figure S2 for more details).

Within each species, cell embeddings for each pattern were separated by MetaMarker cell type and developmental age (Figure 1D). The patterns show cell type specificity and clear dynamics across developmental time as cells transition through the neurogenic trajectory. Especially clear in the detailed time course of the mouse and macaque data are the descent of progenitor patterns p5 and p4of7CtxDev, and the increase in the neuronal maturation pattern, p2, over time. Importantly, these trends are not limited to any one cell type. All classes of neural progenitor have low expression of neuronal patterns p7 and p2, yet show distinct increases in these signals as development progresses. This “neuronalization” of progenitors has been noted previously by Telley 2019 (PMID: 31073041), Polioudakis 2019 (PMID: 31303374), Ji 2023 (PMID: 38464021), and Braun 2024 (PMID: 37824650). Similarly, glutamatergic neurons have low levels of IPC pattern p4 and show further decreases over time. Laser capture microdissection-coupled expression data confirmed these temporal trends in independent macaque and human datasets (Figure S1C). These observations of shared transcriptomic elements across cell types suggest a model of neurogenesis where continuous change coexists with the near binary shift from precursor to post mitotic neuron. In this model, neural progenitors are progressively drawn toward the neuronal transcriptome state, and newborn neurons continue to shut down remaining transcriptomic elements of their precursor state. Emblematic of this, the nascent neuron pattern p7, which is highest in cells considered to be neurons by both the MetaMarker analysis and the original authors, still shows highly significant enrichment in cell cycle genes (Figure 1C). These findings highlight the continuous and overlapping nature of transcriptomic elements resident in individual cells that, especially in development, must be reconciled with the non-overlapping classifications often imposed on single-cells and genes in current multi-omic analyses.

### Exploration of conserved transcriptomic elements of neocortical neurogenesis across NeMO Analytics data collections via transfer learning

To validate and further explore the function of these fundamental molecular programs in neurogenesis, we investigated their expression dynamics across developmental time and space in the NeMO Analytics data collections (Figure 2 & S2A-B). While the joint decomposition in Figure 1B was performed offline, the gene loadings underlying these transcriptomic patterns (or any gene signatures of interest to researchers) can be uploaded to NeMO Analytics (Figure 1A) where transfer learning methods can be used to 1] demonstrate their robustness across measurement technologies, 2] assess their conservation across species and 3] extend our understanding of their temporal and spatial dynamics across development. Projections confirm the cell type mapping of the transcriptomic patterns in an independent scRNA-seq study of the fetal human brain (Figure 2A) and define their laminar (Figure 2B) and single-cell spatial distribution within the classical radial organization of neocortical neurogenesis (Figure 2C). Projection of scRNA-seq of birth-dated RGCs elucidated the temporal progression of neurogenic cells through these transcriptomic programs in the mouse mid-gestational neocortex (Figure 2D): it appears to take ∼1 day for dividing RGCs at the ventricular surface to transition from high p5 (RGCs), through high p4 (IPC), and to high p7 (nascent neuron) transcriptome states. By three days later, neurons are maturing with greatly reduced p7 and high levels of the maturing neuron pattern p2. The notion that p7 is transient while p2 is part of the permanent mature neuron transcriptome is supported by spatial transcriptomic data from the adult human cortex, where p7 shows no systematic expression pattern, while p2 is high across the entirety of the cortical wall and low in white matter (Figure 2F). By directly comparing levels of p7 and p2, this early neuronal transition can be explored in spatial detail in microdissection-coupled expression data from macaque cortex across pre- and post-natal development, where it is evident that as new neurons initiate their radial migration, they also begin their transcriptomic transition from proneural p7 to maturing p2, completing this transition upon arrival at their final destination in the cortex (Figure 2G). This initial maturational transition from p7 to p2 is ubiquitous in neurons of the neocortex and can be seen in new-born neurons across each of the data sets in Figure 2 (Figure S2C-F).

**Figure 2:**
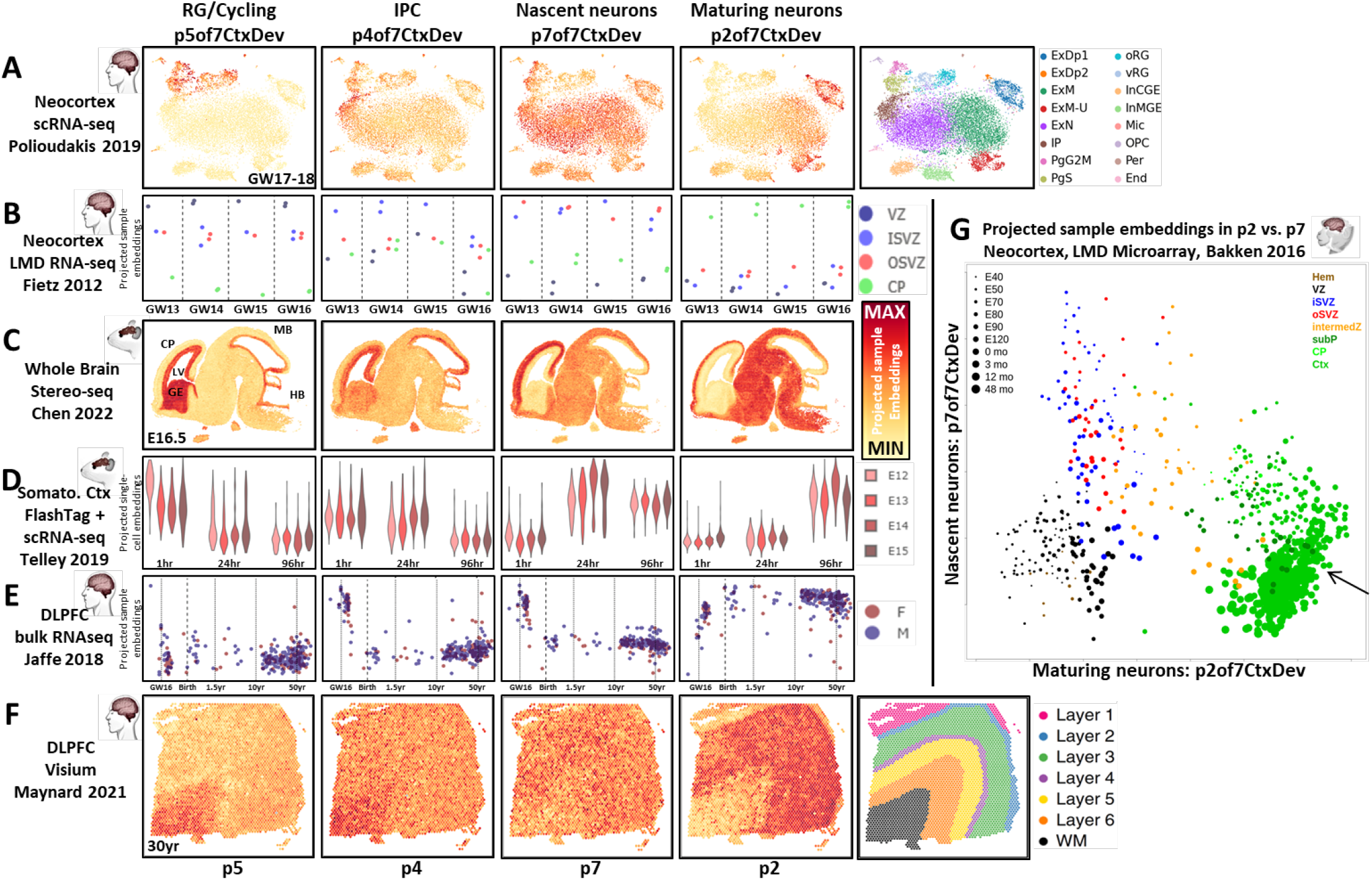
Projection of datasets from the NeMO Analytics collection into transcriptomic dimensions of neocortical neurogenesis yields evolutionary and developmental insights. Each row of panels depicts the projection of a dataset into the p7CtxDev patterns defined in Figure 1; each column is 1 of the 4 highlighted patterns. **A]** tSNE representation of scRNA-seq in fetal human neocortical tissue (PMID: 31303374) colored by the strength of conserved transcriptomic patterns. Original author cell type calls: Ex=exictatory, Dp=deep, N=new/migrating, M=maturing, Ip=intermediate progenitor, Pg=cycling progenitor in S or G2M phase, RG=ventricular (v) or outer (o) radial glia, In=inhibitory neurons of the medial (MGE) or caudal (CGE) ganglionic eminence, Mic=microglia, OPC=oligodendrocyte precursor cell, Per=perictye, End=endothelial cell. **B]** Bulk RNA-seq in laser microdissected (LMD) samples from human fetal neocortex (PMID: 22753484). Y-axis values indicate each sample’s level of the transcriptomic patterns. **C]** Spatial transcriptomics in the fetal mouse brain (PMID: 35512705) colored by levels of each pattern. See Figure S2C for higher resolution comparison of p2 and p7 and Figure S2F for projection across the developmental time course in this dataset. CP=cortical plate, LV=lateral ventricle, GE=ganglionic eminence, MB=midbrain, HB=hindbrain. **D]** scRNA-seq of RGCs labeled at E12-E15 during their terminal division on the ventricular surface at 0hr, then harvested for sequencing at 1hr, 24hr, and 96hr (PMID: 31073041). **E]** Bulk RNA-seq of dorsolateral prefrontal cortical (DLPFC) tissue across the human lifespan (PMID: 30050107). Age is on a transformed log scale to allow better visualization of early development where change is greatest. **F]** Spatial transcriptomics in the adult human dorsolateral prefrontal cortex (PMID: 33558695). **G]** Scatter plot of individually laser microdissected regions of the developing macaque cortex comparing levels of p2 and p7 (PMID: 27409810). Hem=cortical hem, VZ=ventricular zone, ISVZ=inner subventricular zone, OSVZ=outer subventricular zone, intermedZ=intermediate zone, subP=subplate, Ctx=Cortex. Arrow indicates mature neurons of the cortex, where p7 has descended and p2 is highest. See Figure S2A-D for this p2 vs. p7 analysis in additional datasets. With the exception additional labels and panel G, this entire figure was created from NeMO Analytics screen captures. Units resulting from projection analyses are comparable only within, not across, projected datasets. For this reason, in this report we display all data projections on a minimum to maximum scale bounded by each individual dataset projected (Methods). Expression of individual genes can be explored in these specific datasets at NeMOlink05 and the 7 jointNMF transcriptomic patterns (p7CtxDev) at NeMOlink06.

Interrogating the collection of neocortical development data with transfer learning tools in NeMO Analytics yielded broad evolutionary and developmental insights into these jointly defined transcriptomic dimensions. They can be further explored across many additional *in vivo* single-cell and spatial datasets through projection analysis at: NeMOlink07. We invite researchers to explore these and add their own emerging datasets and gene lists of their own interest to expand the discovery potential of this centralized public neocortical development resource (tutorials at carlocolantuoni.org).

### Higher resolution dissection of the conserved neurogenic transcriptome in neocortex yields insight into oRG evolution

To dissect this same sub-collection of neocortical data at higher resolution, we performed a second jointNMF decomposition of the same 3 scRNA-seq data sets from mid-gestation, in this case defining 40 dimensions (patterns from this decomposition will be referred to using the “p40CtxDevo” suffix). This analysis yielded more detailed transcriptome dynamics across both progenitors and neurons of the neocortex along with more cell type specific genetic associations (Figure S3A and Table S2). Of particular interest in this analysis was a pattern expressed at high levels in thousands of RGCs of the macaque and human, but only in a small number of RGCs in the mouse (Figure 3A, p27of40CtxDev). These cell populations coincide with outer radial glia (oRG or basal RG, bRG) cell type calls by original authors in the macaque and human studies (Figure S3B). Additionally, the ranking of the human oRG markers HOPX, FAM107A, MOXD1, TNC and the ligand-receptor pair PTN-PTPRZ1 (PMID: 26406371) were all in the top 100 genome-wide loadings for this pattern (Figure S3 and Table S2), indicating that p27of40CtxDev represents a transcriptomic program in oRG cells. oRG cells are a primate and human expanded cell type linked to evolutionary increases in neuron number, cortical surface area, and gyrification (PMID: 20436478, PMID: 21127018). Consistent with our observation of the sparsity of high p27 signals in single cells of the mouse neocortex, oRG cells account for 40-75% of dividing cells in the developing human neocortex (PMID: 20154730), while accounting for <10% in the mouse (PMID: 21478886).

**Figure 3:**
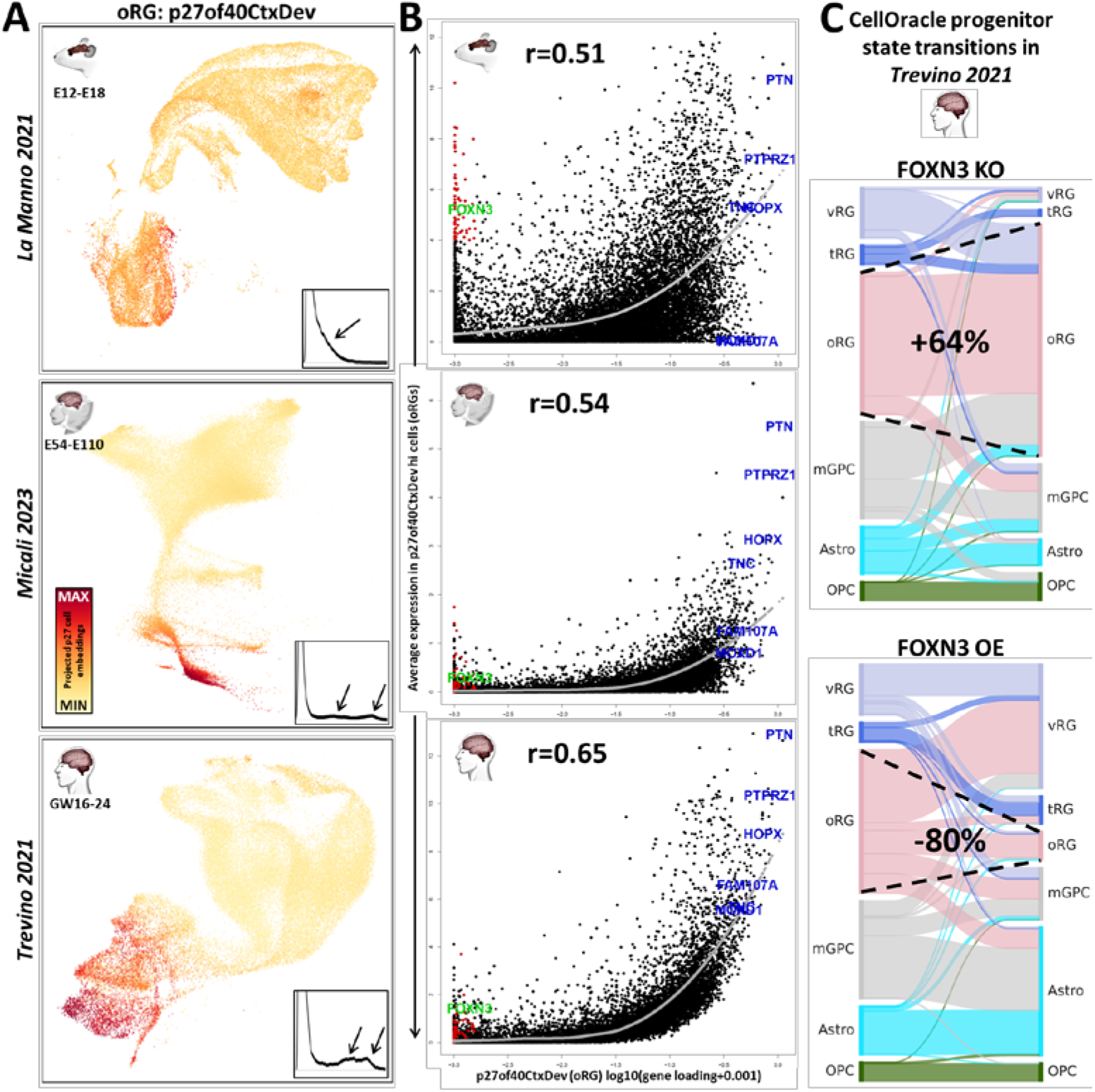
Higher resolution decomposition of the developing neocortical transcriptome yields insight into oRG evolution. **A]** One of 40 patterns (**p40CtxDev**) defined in a higher resolution jointNMF decomposition of neocortical development across mouse, macaque and human: pattern p27of40CtxDev is a partially conserved transcriptomic elements of the oRG cell type across mammalian species. Single-cell embeddings for p27 are shown in a color gradient across the low-dimensional representation of cells in all 3 species. Inset plots show the distribution of p27 embeddings in each species. Arrows indicate largest deviations between mouse and NHP & human distributions. Table S2 contains gene loadings for the entire set of 40 transcriptomic patterns along with their enrichments in disease and cell biological gene lists. **B]** Scatter plots of p27 gene loadings against average expression in oRG cells (defined by levels of p27) in each species. oRG marker genes are shown in blue. Genes in red have low loadings in p27, but have high expression in putative mouse oRG cells - among these, FOXN3 is shown in green. The gray curve is a loess fit of the average expression of genes across the magnitude of gene loadings in this pattern. Correlations of these 2 measures are noted in each species (p<2.2e-16 in each case). **C]** Cell type transitions predicted by *in silico* FOXN3 knock-out (KO) and over-expression (OE) simulations in a CellOracle analysis which integrated scRNA-seq and scATAC-seq data from neural progenitors in Trevino 2021 (PMID: 36755098) to construct regulatory networks in the developing neocortex. Dashed lines show the expansion of the oRG cell type in FOXN3 KO and its reduction in FOXN3 OE. Images in panel A were created from NeMO Analytics screen captures. All 40 patterns can be explored across mammalian neocortical developmental data at NeMOlink08. vRG=ventricular radial glia, oRG=outer radial glia, tRG=truncated radial glia, mGPC=multipotent glial precursor cells, Astro=astrocytes, OPC=oligodendrocyte progenitor cells.

To further explore the individual genes involved in this partially conserved transcriptomic signature of oRGs, in Figure 3B we have plotted each gene’s loading in p27of40CtxDev against the average expression of that gene in the mouse, macaque or human cells that have highest levels of this oRG signature (Methods). In each of the 3 species, we observed the expected positive correlation between the p27 loadings and expression levels in oRG cells high in p27. Consistent with a cell type of evolutionarily increasing cohesiveness, this correlation grows from mouse to macaque to human, with the relative expression of canonical markers of human oRG cells increasing along this same evolutionary trajectory (Figure 3B). This suggests a model in which a transcriptomic regulatory program that began as a diffuse network in progenitor cells of the rodent-primate common ancestor has evolved over time to drive the emergence of a novel progenitor type central in the expansion of the neocortex in the primate and human lineages. This is consistent with our recent observations employing independent methods, which indicate that the first evolutionarily components of the oRG transcriptomic program arose in gliogenic precursors of the rodent-primate ancestor before expanding and driving the evolution of oRG cells in the primate lineage (PMID: 37383947; Figure S3C).

In the mouse specifically, there are many genes that have near-zero loadings in p27of40CtxDev (i.e. genes NOT involved in the p27 transcriptomic element of oRGs) while still having high expression in the cells with the highest p27 levels in the developing murine neocortex (Figure 3B, red highlighted genes at low X-axis values). In contrast, in both the macaque and human oRG cells, no genes with low p27 loadings are expressed at high levels. This raises the possibility that these genes are part of a transcriptomic program that has been shut down in oRG cells since the rodent-primate divergence. The transcriptional repressor FOXN3 is one of the genes that is low in p27 gene loadings and high in mouse oRG cells but not in the macaque or human cells (Figure 3B, in green). Strikingly, FOXN3 target genes are enriched among high p27 gene loadings (targets from MsigDB; p=3.9e-20). To explore this further, we integrated scRNA-seq and scATAC-seq data from the Trevino human dataset (PMID: 34390642) in CellOracle (PMID: 36755098) where it is possible to test the simulated effects of TF perturbations on cell identity *in silico*. We found that FOXN3 knock-out (KO) simulation produced many cell state transitions to the oRG state (increasing oRG numbers by 64%), while over-expression (OE) lead to its near disappearance (decreasing oRG numbers by 80%; Figure 3C). This suggests the possibility that high FOXN3 expression in mouse progenitor cells destabilizes the oRG state, reducing numbers of the mouse oRGs, consistent with their sparsity in the rodent brain. This data and the role of FOXN3 as a transcriptional repressor and proliferation inhibitor (PMID: 12808094; PMID: 24403608; PMID: 27259277) are consistent with a model in which primate oRG cells, where FOXN3 expression is low, allow the increased expression of FOXN3 targets, resulting in the observed enrichment of these targets in p27 gene weights, and the stabilization of the proliferative oRG cell state in primate and human neocortex. Hence, it appears that the evolution of the oRG cell type in primates may have involved both subtraction from and addition to the ancestral rodent RGC transcriptomic program.

### Joint decomposition defines excitatory neuronal laminar identities in the adult neocortex

To generate a precise molecular definition of mature layer-specific neuronal transcriptome identities, we interrogated single-nucleus RNA-seq (snRNA-seq) data from adult human neocortical tissue from Jorstad 2023 (PMID: 37824655) and Bakken 2021 (PMID: 34616062). Again using the jointNMF approach in SJD (doi.org/10.1101/2022.11.07.515489), we included only Smart-seq v4 data (for more complete transcript coverage and deeper sequencing) from excitatory neurons derived from layer-microdissected tissue. This joint decomposition defined 20 neuronal transcriptome signatures that are shared across 5 snRNA-seq data matrices, each from a distinct donor spanning a total of 8 neocortical regions in these two studies (Figure S4). We confirmed the layer-specific distribution of these mature human neuronal patterns across species and measurement technologies using transfer learning methods in NeMO Analytics: Figure 4 shows the projection of spatial transcriptomic and additional snRNA-seq data from adult human, macaque and mouse neocortex into 9 of the 20 adult human neuronal transcriptome patterns (patterns from this decomposition will be referred to using the “p20CtxLayer” suffix) some of which shared transcriptome signatures with specific subcortical neuronal identities (Figure 4D & S4).

**Figure 4:**
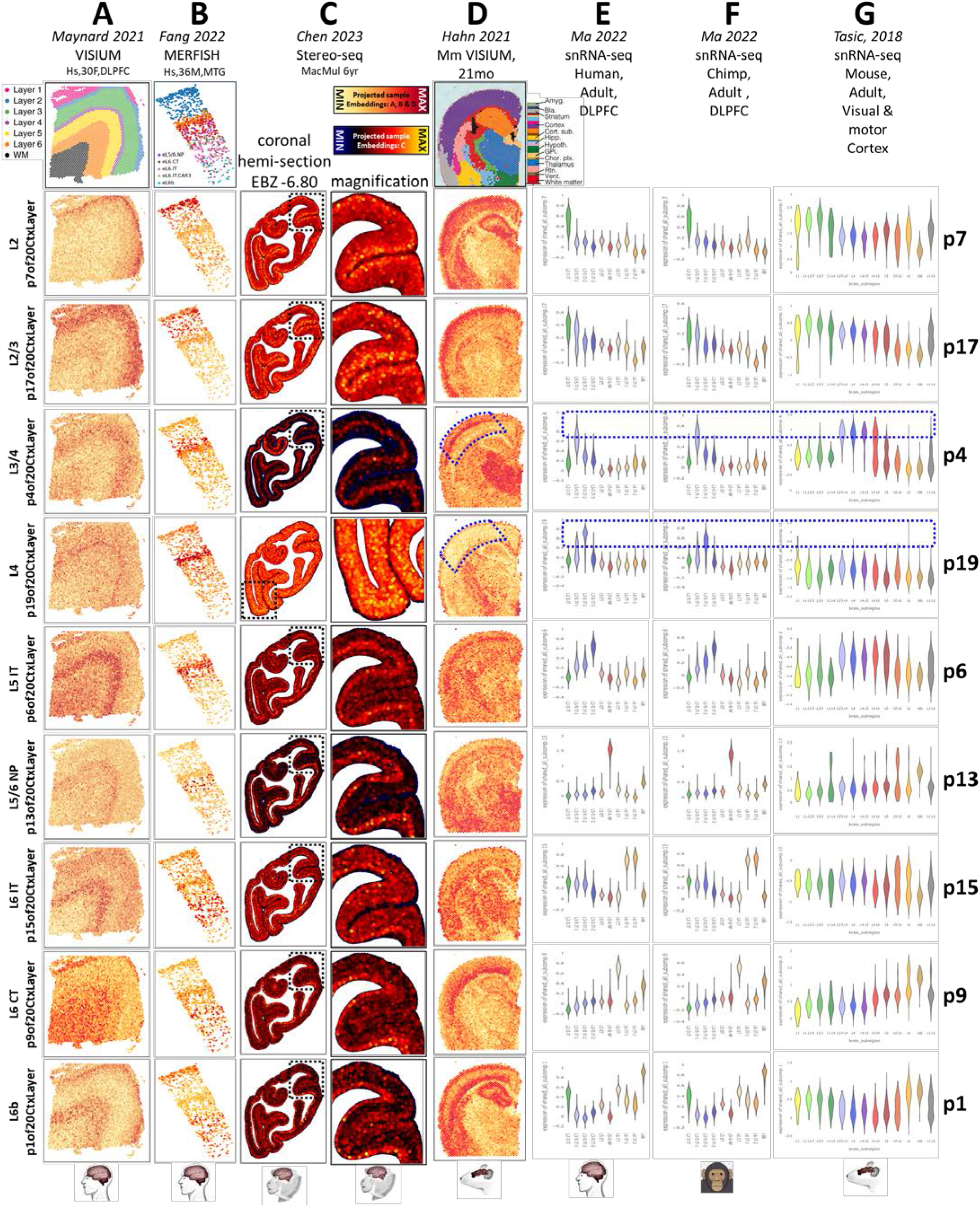
Mature human neocortical layer-specific neuronal transcriptome signatures across mammalian species. Each column shows the projection of one dataset into 9 of the 20 layer-specific signatures (**p20CtxLayer**). Each row thus depicts the expression level of a signature across spatial **A-D** and single-cell **E-G** transcriptomic datasets from adult human, primate and mouse (PMID: 33558695, PMID: 35771910 PMID: 37442136, PMID: 37591239, PMID: 36007006, PMID: 30382198). Original author cell type calls were used. CT=corticothalamic, IT=intratelencephalic, NP=near projecting. Blue dashed annotations indicate layer 4 patterns which are conserved across all three mammals (p4) or which are primate-specific (p19). Due to heterogeneity in the p19 signal, we have magnified a different region to show this pattern in panel C. It is unclear if this heterogeneity is due to regional specificity or signal to noise variation. With the exception of additional labels, this entire figure was created from NeMO Analytics screen captures. An expansive collection of adult neocortical data at NeMO Analytics can be explored using individual genes (NeMOlink09) or these jointNMF patterns (NeMOlink10). Table S3 contains gene loadings and the full gene set enrichments across all 20 patterns.

Pattern p4of20CtxLayer identifies a layer 4 neuronal identity that is conserved in the neocortex of all 3 mammalian species, while pattern p19 marks a distinct layer 4 transcriptomic identity that can be seen in both human and macaque, but not mouse neocortex (Figure 4, blue dashed annotations in p4 and p19). Ma 2022 (PMID: 36007006) recently observed primate specific expression of FOXP2 (a gene that has been implicated in human language development and neuropsychiatric disease; PMID: 11586359; PMID: 12687690) in excitatory neurons of layer 4. FOXP2 is ranked in the top 1% of genome-wide loadings for the primate-specific layer 4 pattern p19. Similarly, Chen 2023 (PMID: 37442136) reported several primate specific layer 4 neuronal cell types in a recent study of spatial transcriptomics in the adult macaque brain. Interrogation of the genes reported in the Chen layer 4 signal in p19 gene loadings revealed a significant enrichment of high values in this small group of 11 genes (p=4.5e-6), which again included FOXP2 (Figure S4). Hence, the primate-specific layer 4 pattern, p19, that we have defined here is likely detecting the same FOXP2-related layer 4 signal as that described in Chen 2023 (PMID: 37442136) and Ma 2022 (PMID: 36007006).

We observed that layer 4 transcriptomic identities are present in cells of agranular cortical regions, e.g. primary motor cortex (Figure S4), as seen in Jorstad 2023 (PMID: 37824655). This reinforces the notion that although some cortical regions lack histologically defined layer 4 pyramidal cells, shared transcriptomic identities are indeed present in morphologically distinct cells in approximately the same laminar position. Notably, this p19of20CtxLayer transcriptomic program is also primate-specific when observed in non-layer 4 neurons (Figure S4). Similar to the observation of the oRG transcriptomic program examined in Figure 3, this suggests that new species-specific cell types may arise through the evolution of gene regulatory programs that are initially present in conserved cell types. This is consistent with our understanding that regulatory variation is much more wide-spread than cell type or protein coding variation, both across species and within the human lineage.

### Mapping the developmental emergence of adult neuronal laminar identities in the neocortex

We next sought to explore the developmental emergence of these layer-specific neuronal elements of the adult human neocortical transcriptome. Employing transfer learning methods implemented in Nemo Analytics, we explored their expression in snRNA-seq data from neurons in fetal, postnatal, and mature human neocortex from Herring 2022 (PMID: 36318921; Figure 5A), and in laser capture microdissection (LMD)-coupled microarray data spanning pre- and post-natal macaque neocortical development from Bakken 2016 (PMID: 27409810; Figure S5A). When visualized on the UMAP of the snRNA-seq data, each of the patterns appears to occupy a distinct laminar-specific neuronal identity spanning fetal and postnatal development (Figure 5A & S5C), suggesting that the mature laminar identities begin to emerge early in development. To confirm this, we depicted these same projections as strip charts across age (showing expression levels in individual cells: Figure 5B) and line plots across age (to summarize lifespan trends: Figure S5B) for each of the neuronal subtypes defined in Herring 2022. Each of the adult layer patterns is more highly expressed in one of the specific neuronal subtypes distinguished in the fetal data, and all of the neuronal laminar identities build over developmental time, with lowest levels during fetal ages and increasing expression over many years of postnatal life. This protracted timeline for the full acquisition of lamina-specific identities is similar to many elements of human brain development that have acquired longer periods of maturation over evolutionary time (PMID: 30545855; PMID: 37003107).

**Figure 5:**
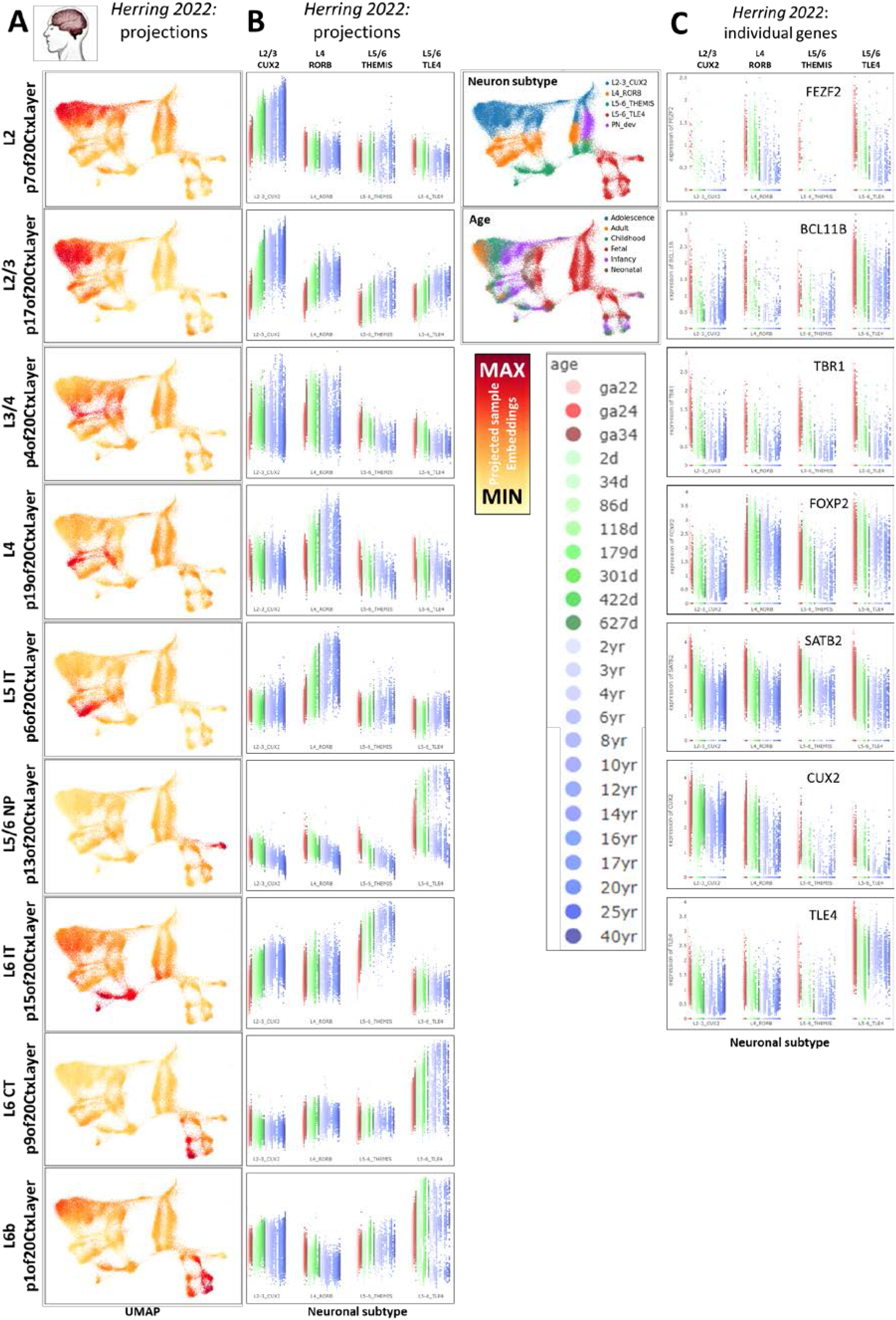
Projection of fetal and postnatal neuronal snRNA-seq data into adult layer-specific neuronal transcriptome patterns (p20CtxLayer from Figure 4). **A]** Projection of neuronal data from Herring 2022 (PMID: 36318921) into the p20CtxLayer patterns, displayed as color scales in UMAP dimensions and **B]** as strip charts with individual cell embeddings across cell types (defined by original authors) and ages. See Figure S5A for laminar specificity and maturation timing in the macaque. **C]** Many conventional neuronal TF marker genes for specific cortical layers peak at the earliest fetal time points observed here. With the exception of additional labels, this entire figure was created from NeMO Analytics screen captures. These and additional detailed visualizations of the p20CtxLayer jointNMF patterns across neocortical datasets can be explored in Figure S5, and specifically in the Herring 2022 data in Figure S5 and at NeMOlink11.

This prolonged maturation of neuronal laminar transcriptomic identities is in stark contrast to the timing of expression of the individual transcription factor (TF) genes often employed to distinguish neurons of different neocortical layers (Figure 5C). Many of these TFs show highest and most specific expression at early fetal time points, after which their RNA levels descend as the adult laminar identities that they drive continue to build and take years to reach full maturity. A specific example of this general phenomenon has recently been examined in great detail in Nano 2023 (PMID: 37745597), where the neuronal lineage-defining TF, FEZF2, was shown to reach peak expression at early fetal ages, just as the deep layer transcriptomic identity which it constructs is beginning to emerge. This suggests that neuronal identity defining TFs setup up lasting epigenetic structure during fetal development, enabling the stable execution of layer specific transcriptional programs through years of maturity in the absence of high levels of their own mRNA. Hence, while canonical TF markers of neuronal laminar-specific identities are very effective for the determination of neuronal subtypes in early prenatal development, they are likely poor metrics of layer-specific neuronal maturation, for which our transcriptome-wide jointNMF patterns from adult neocortical data are ideal (we employ them in this capacity below in Figures 6 and 8).

**Figure 6:**
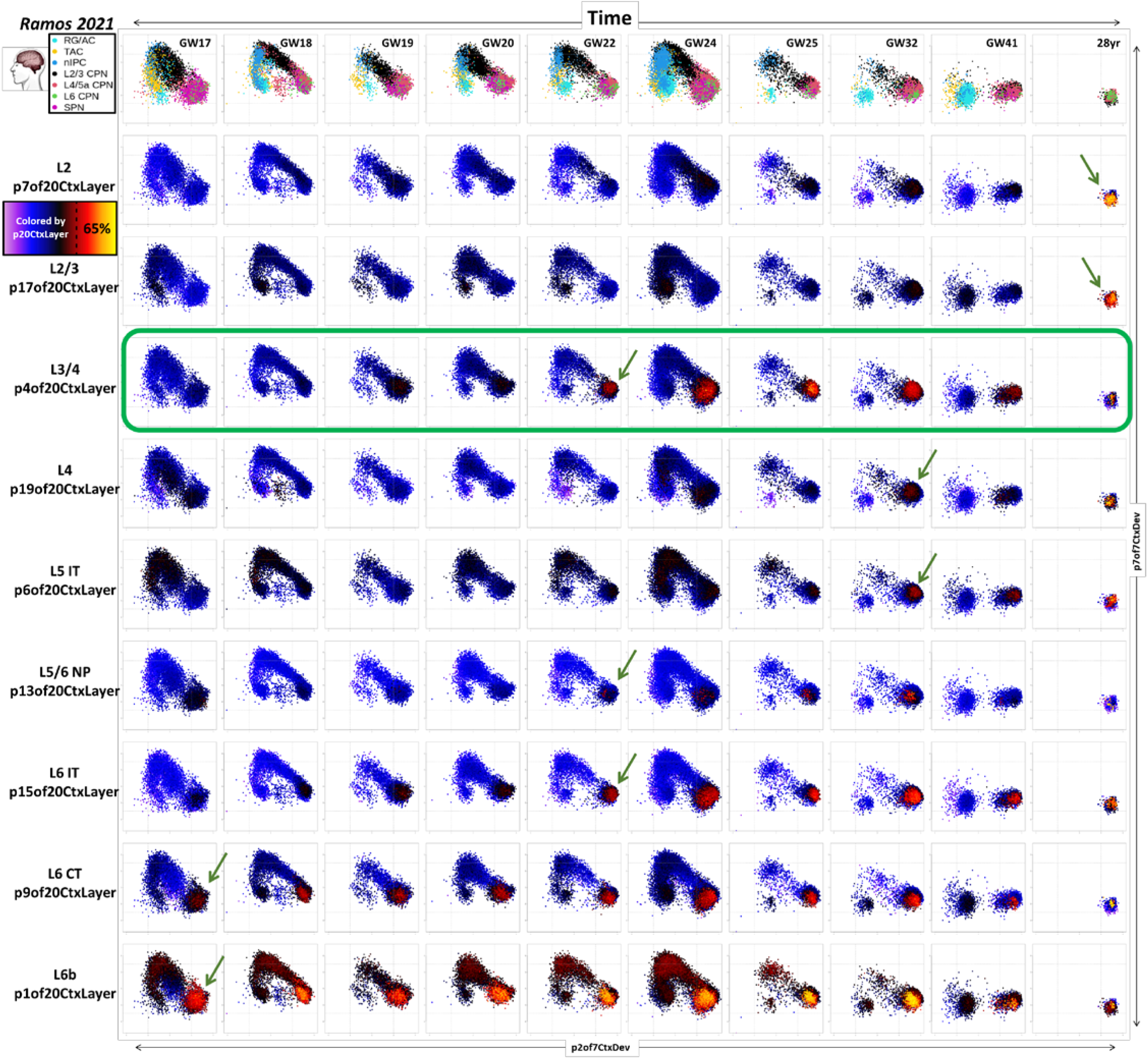
Mapping neuronal maturation and the emergence of specific laminar identities across development of the human neocortex. Plots represent projection of snRNA-seq data from pre- and post-natal human neocortex (Ramos 2021, PMID: 36509746, including only cells in the neocortical excitatory neurogenic lineage) into transcriptomic dimensions that define neuronal birth and maturation. Each column shows data from a single donor. The X-axis in each plot maps the individual cells onto p2of7CtxDev (neuron maturation). The Y-axis maps cells onto p7of7CtxDev (proneural/nascent neurons). The color of points in each row shows the strength of one of the 9 transcriptomic programs defined in adult layer-specific neuronal data (p20CtxLayer, see Figure 4 & 5). Green arrows indicate the earliest age at which cells surpass 65% of the maximal level for each signature (Figure S6 for details). The L3/4 p4of20CtxLayer pattern emerges earlier than other upper layer neocortical neuronal identities (green box). Original author cell type calls are used. RG/AC=radial glia / astrocytes, TAC=transit amplifying cells, nIPC=neuronal intermediate progenitor cells, CPN=cortical projection neurons of different layers, SPN=subplate neurons. Additional *in vivo* data spanning early postnatal ages are explored in this manner in Figure S6.

To chart the initial appearance and extended maturational trajectories of specific cortical laminar identities, we combined the broad elements of neuronal maturation captured in the low-resolution cortical transcriptome decomposition (Figures 1&2; p7CtxDev) with the adult layer-specific neuronal transcriptomic programs (Figures 4&5; p20CtxLayer): Figure 6 shows the projection of human snRNA-seq data from pre- and post-natal time points (Ramos 2021, PMID: 36509746) into the two sequential dimensions of new-born neuronal development: p7of7CtxDev (nascent neurons) and p2of7CtxDev (neuron maturation). As in Figures 2G & S2C-F, this 2D space describes pan-neuronal birth and early maturation where neurons are born from progenitors at the low p7 & low p2 state, transiently express the proneural p7, and finally repress p7 and induce p2 as they arrive at their ultimate position in the developing neocortex. At ages prior to and including GW24 this full arc can be seen in Figure 6. After GW24, as progenitors and nascent neurons begin to disappear, high p7 and low p2 states are vacated and cells coalesce into a single population of maturing neurons at the low p7 & high p2 state. This space alone does not distinguish neuronal subtypes, hence, cells in each row of plots across age in Figure 6 are colored by levels of one of the adult neuronal patterns (p20CtxLayer). These plots demonstrate that the laminar-specific transcriptomic identities begin to emerge only after the nascent neuron pattern p7 has been shut down and the maturing neuron pattern p2 has been maximally induced, i.e. as new neurons arrive at their final laminar destination. This suggests that induction of neocortical neuronal laminar identities requires the cellular environment specific to laminar position in the cortex and are not fully intrinsic to neurons born at a particular time or place.

In general, the time at which each laminar-specific pattern begins to appear follows the classic inside-out (deep-to-upper) developmental architecture of the neocortex (Figure 6, green arrows). p1of20CtxLayer (subplate & Layer 6b) appears to emerge even before the times sampled in this data, peaking between GW22-32 and then descending slightly (likely as the subplate disappears while layer 6b neurons persist). While it is clear that layer 2/3 patterns p17 and p7 reach peak levels postnatally, the time points studied in Ramos 2021 (PMID: 36509746) do not span early postnatal life. Data in Herring 2022 (PMID: 36318921) indicate that these patterns appear strongly in the months following birth and largely plateau after 2 years (Figure 5A, S5 & S6). The conserved mammalian layer 3/4 pattern p4 is an exception to the classical sequence of layer-specific neuronal appearance, emerging earlier than other adjacent neuronal laminar identities, which can also be seen in the Herring 2022 data (Figure 5A&B, S5B, & S6). These observations link to recent findings from Huilgol 2024 (PMID: 38645016) who observed that in the developing mouse neocortex particular layer 3/4 neurons are born days before the majority of other neurons in these layers. We see a similar deviation from the strict inside-out laminar developmental progression in the Ramos 2021 (PMID: 36509746) human data where neurons with high p4 levels appear weeks prior to other neurons in the same laminar position. Further investigation is required to reveal the exact identity of these precociously maturing neurons and determine if they are the same as the precociously born neurons observed in Huilgol 2024 (PMID: 38645016). We present this approach of mapping novel data into well-characterized dimensions of cellular dynamics from previous datasets as a general approach to leverage increasing amounts of new data in charting precise elements of development. Similarly, we next apply this methodology to explore specific elements of transcriptomic change from *in vivo* development which are paralleled in stem cell derived neural differentiation systems.

### Broad in vivo transcriptome dynamics are recapitulated in vitro, while specific mature neuronal laminar identities are incomplete

Human pluripotent stem cell (hPSC)-derived models have become central tools in modeling neocortical development and disease. Understanding which elements of development are and are not modeled with high fidelity is essential to using these systems effectively to develop novel therapeutics (PMID: 38915580). We used the transcriptomic dimensions that we defined within *in vivo* neocortical neurogenesis and maturation to interrogate *in vitro* models of cortical development. First, we projected several *in vitro* datasets into transcriptomic patterns from Figure 1 & 2 (p5, p4, p7, and p2of7CtxDevo). The sequential progression through the broad elements of *in vivo* neocortical neurogenesis defined in these patterns is clear in bulk RNA-seq from 2-dimensional differentiation, scRNA-seq from organoid models, and spatial transcriptomic data in organoids (Figure 7 & S7). These same datasets were also projected into the transcriptomic signature of oRG (27of40CtxDev from Figures 3 & S3) showing that the early precursor pattern p5of7CtxDev, which is high in dividing RG *in vivo*, appears *in vitro* well before p27of40CtxDev (arrows in Figure 7 & S7).

**Figure 7:**
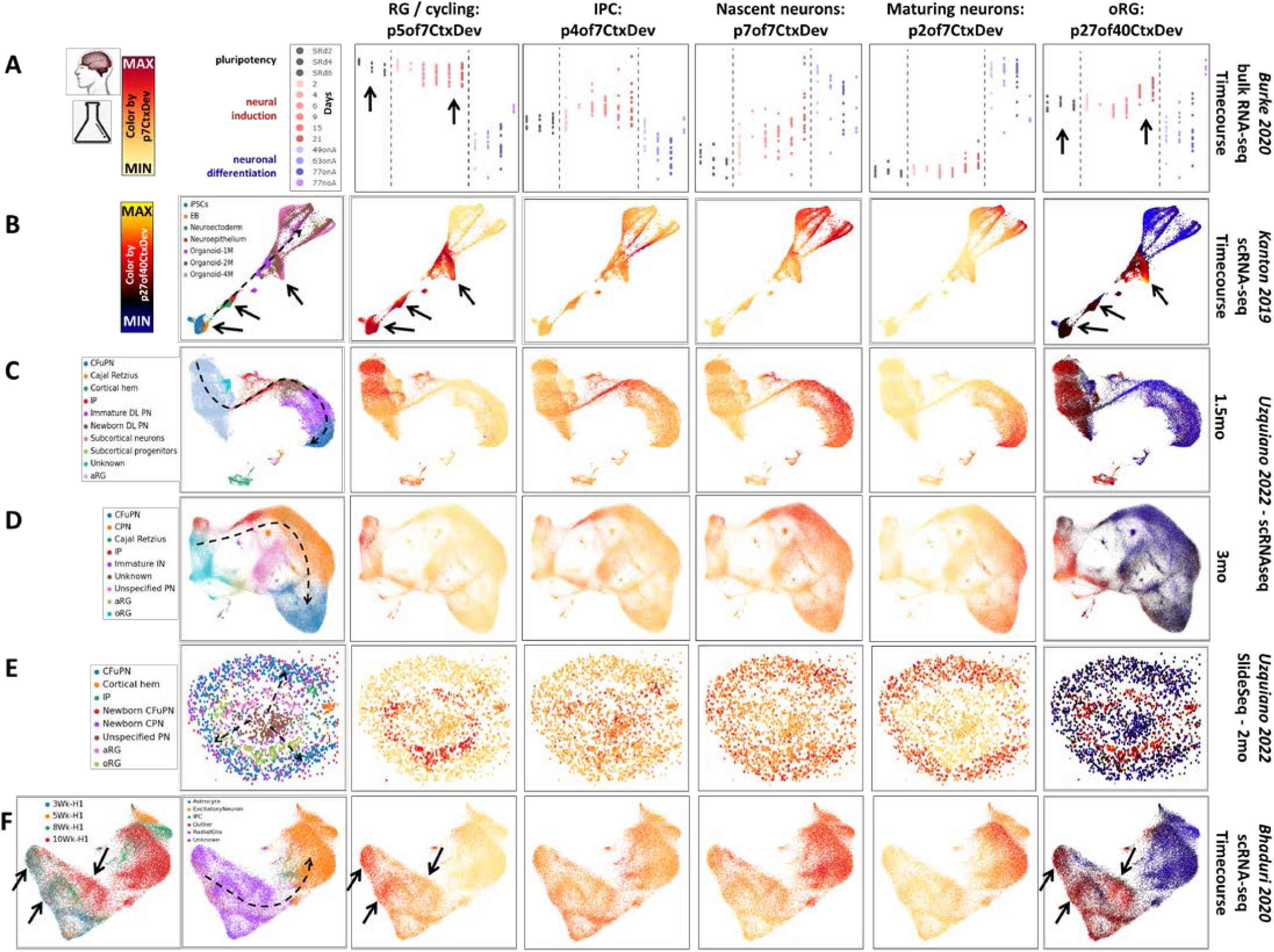
Broad elements of in vivo development are recapitulated in vitro: Projection of data from *in vitro* neural differentiation models into the p7CtxDev patterns from Figure 1 and p27of40CtxDev from Figures 2&3. Projection of the oRG transcriptomic signature, p27of40CtxDev, is shown in a different color scale, indicating that it was derived from a distinct joint decomposition than the other patterns. **A]** Bulk RNA-seq data from 2D *in vitro* differentiation of 14 hPSC lines from 5 donors. SRd=days in pluripotent self-renewal, days of neural induction in red, neuronal differentiation in blue/purple, onA=astrocyte co-culture, noA=no astrocytes in culture (PMID: 31974374). **B]** scRNA-seq data from pluripotency through 4 month cerebral organoids (PMID: 31619793) in a force-directed graph layout. **C&D]** scRNA-seq at single time points in cerebral organoid differentiation (PMID: 36179669) in UMAP plots. **E]** Spatial transcriptomics in a 2 month cerebral organoid (PMID: 36179669). **F]** scRNA-seq across 3-10 weeks of cerebral organoid differentiation in a single hPSC line using the “more directed” Xiang 2017 (PMID: 28757360) protocol from (PMID: 31996853) in a UMAP plot. Dashed lines indicate approximate neurogenic trajectories in each experiment. This *in vitro* transfer learning experiment examining broad elements of neurogenesis (p7CtxDev) parallels that performed in Figure 2 where *in vivo* data was used. Original author cell type calls were used: EB=embroid body, Cfu=corticofugal, PN=projection neurons, DL=deep layer, IN=inhibitory neuron, aRG=apical radial glia, oRG=outer radial glia, IP=intermediate progenitor. With the exception additional labels, this entire figure was created from NeMO Analytics screen captures. These transcriptomic patterns can be explored across these data and a larger collection of *in vitro* differentiation data at NeMOlink12 and NeMOlink13 and individual genes at NeMOlink14. Arrows indicate time points at which the expression of p5of7CtxDev and p27of40CtxDev differ - this is explored in more depth in Figure S7.

Notably, while *in vivo* neuronal data clearly show reductions in p7of7CtxDev (nascent neuron) in cells where p2of7CtxDev (neuron maturation) peaks (Figures 2, S2, & 6), this is not as clear in the *in vitro* models (Figure 7). To examine the neuronal maturation of these systems in more depth, we recreated the transfer learning experiment performed on *in vivo* data in Figure 6, here employing *in vitro* cerebral organoid data (PMID: 31619793, PMID: 36224417; Figure 8A & S8).

**Figure 8:**
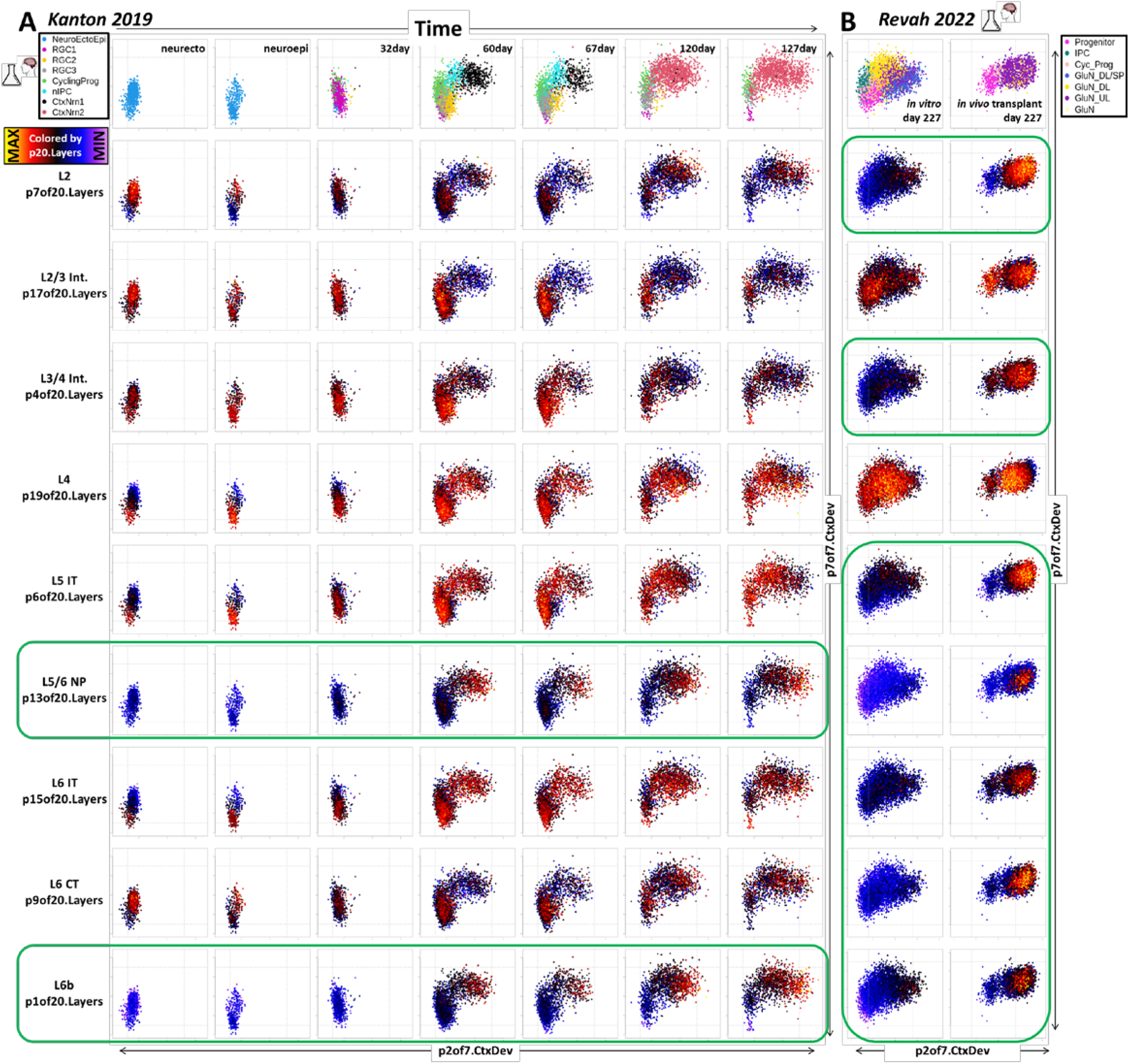
Mapping neuronal maturation and the emergence of specific laminar identities in hPSC-derived models of neocortical neurogenesis. **A]** Projection of scRNA-seq data from an *in vitro* cerebral organoid time course (PMID: 31619793, including only cells in the neocortical excitatory neurogenic lineage) into transcriptomic dimensions that define neuronal birth and maturation. Each column shows data from a single time point. The X-axis in each plot maps the individual cells onto p2of7CtxDev (neuron maturation). The Y-axis maps cells onto p7or7CtxDev (proneural/nascent neurons). The color of points in each row shows the strength of one of the 9 transcriptomic programs defined in adult layer-specific neuronal data (p20CtxLayer). Green boxes indicate where specific laminar identities (p1:L6b and p13:L5/6 NP) follow trajectories similar to *in vivo* development, i.e. absent from early progenitors at low p7 + low p2 levels and appearing systematically in neurons at high p2 levels following a rise and fall of p7. While only data from 1 hPSC line is shown here (409b2), the 2^nd^ line used in the Kanton 2019 (PMID: 31619793) time course study (H9) showed the specific maturation of these same 2 neuronal identities, as do additional studies (Figure S8). **B]** Transplantation of human cerebral organoids into the cortex of newborn rat pups elicited significant additional neuronal maturation along specific laminar trajectories over conventionally grown organoids (PMID: 36224417). Green boxes indicate where specific laminar identities follow trajectories similar to *in vivo* development. Transplantation appears to increase emergence of all but 2 of the layer specific maturational signatures. Paradoxically, while neurons in transplanted organoids showed much elevated levels of specific neuronal identities and p2 over their *in vitro* counterparts, they did not show more reduction of p7. This *in vitro* transfer learning experiment parallels that performed in Figure 6 that used *in vivo* data. Original author cell type calls were used: NeuroEctoEpi=neurectodermal and neuroepithelial states, RGC=radial glial cells, CyclingPrg=cycling neural progenitors, nIPC=neuronal intermediate progenitor cell, CtxNrn=cortical neuron.

Early time points contain primarily progenitor states at low p7of7CtxDev and low p2of7CtxDev states. As neurogenesis begins, the elevation of proneural pattern p7, followed by maturation pattern p2 is clear. Further, as p2 continues to rise, p7 begins to fall. These dynamics parallel those defined *in vivo*, however, neurons *in vitro* fail to complete this maturational trajectory. At later time points, as progenitors and early neurons disappear from low p7 and low p2 states, neurons do not continue to reduce p7 and increase p2, failing to arrive together at a unitary low p7 & high p2 state where mature neuron laminar identities emerge as they do *in vivo* (Figure 6). Importantly, peak mature laminar signals are scattered across cell types and are systematic within the appropriate low p7 & high p2 neurons only in the subplate/L6b pattern p1of20CtxLayer and the L5/6 NP pattern p13, indicating that only these specific laminar neuronal identities are progressing beyond the very earliest stages of maturation in the organoids (Figure 8A & S8, in green).

We have also interrogated the neuronal laminar identities (p20CtxLayer, from Figures 4, 5, & 6) in data from cerebral organoids generated using diverse protocol modifications, including different iPSC lines (PMID: 31619793; Figure S8), more or less regionally directed differentiation (PMID: 34616070), longer time courses (PMID: 33619405; Figure S8), and slice preparations (PMID: 30886407). Each of these experiments showed impact on neuronal maturation trajectories, but none fully recapitulate *in vivo* development along the dimensions that we have defined here (NeMOlink15). While difficult to fully assess without a more detailed time course, transplantation of cerebral organoids into the cortex of new born rat pups (PMID: 36224417) induced increased levels of nearly all the adult laminar signatures over organoids cultured continuously *in vitro* (Figure 8B, in green). Exceptions to this included the L2/3 (p17of20CtxLayer) and the primate-specific L4 (p19of20CtxLayer) patterns, which show high levels that are not systematically localized to the most mature neurons (Figure 8B, not in green). While this clearly indicates that the in vivo environment supports more complete neuronal maturation, it is also an indication of the limitations of the rodent cortex in inducing primate-specific elements of neocortical maturation. Continued interrogation of cerebral organoid data as protocols evolve will be necessary to continually asses what elements of *in vivo* development can effectively be explored *in vitro*.

## Discussion

NeMO Analytics data resources and our joint decomposition of transcriptomic dynamics in neurogenesis and maturation can be leveraged to explore neocortical development and in the design of manipulations of precise cellular mechanisms underlying risk for common complex brain disorders in tractable *in vitro* systems. We invite the research community to explore this collection of public data resources along with the transcriptomic elements of the human neocortex that we have defined and their transfer into *in vitro* stem cell models at nemoanalytics.org (nemoanalytics.org/landing/neocortex). Researchers are welcome to upload their own datasets and gene signatures for dissemination and exploration in this neocortical development research environment. It is our hope that the collective exploratory and communication benefits of housing data in this shared environment will incentivize deposition of emerging neocortical data and data-driven scientific interaction. We suggest researchers complement their deposition of newly published data in traditional raw data repositories with upload to NeMO Analytics where it will be immediately available to researchers with and without coding experience for exploration alongside the compendium of data resident in NeMO analytics (carlocolantuoni.org). We propose these analyses as specific applications of the general approach of combining joint decomposition with large curated collections of analysis-ready multi-omics data matrices focused on particular cell and disease contexts.

## NeMO Analytics web links listed in the manuscript

NeMO Analytics Neocortex Development landing page and guide: https://nemoanalytics.org/landing/neocortex

NeMOlink01: https://nemoanalytics.org/p?l=NeocortexDevoHsInVivo&g=EOMES

NeMOlink02: http://nemoanalytics.org/p?p=p&l=NeocortexDevoHsInVitro&c=CellCycleGenes.Seurat&algo=pca

NeMOlink03: https://nemoanalytics.org/p?p=p&l=Sonthalia2024fig1&c=MammCtxDev.jNMF.p7&algo=nmf

NeMOlink04: https://nemoanalytics.org/p?l=Sonthalia2024fig1&g=EOMES

NeMOlink05: https://nemoanalytics.org/p?l=Sonthalia2024fig2&g=MYT1L

NeMOlink06: https://nemoanalytics.org/p?p=p&l=Sonthalia2024fig2&c=MammCtxDev.jNMF.p7&algo=nmf

NeMOlink07: https://nemoanalytics.org/p?p=p&l=NeocortexEvoDevo&c=MammCtxDev.jNMF.p7&algo=nmf

NeMOlink08: https://nemoanalytics.org/p?p=p&l=NeocortexEvoDevo&c=MammCtxDev.jNMF.p40&algo=nmf

NeMOlink09: https://nemoanalytics.org/p?&l=AdultNeoctxLayers&g=FOXP2

NeMOlink10: https://nemoanalytics.org/p?p=p&l=AdultNeoctxLayers&c=HsCtxLayer.jNMF.p20&algo=nmf

NeMOlink11: https://nemoanalytics.org/p?p=p&l=Herring2022&c=HsCtxLayer.jNMF.p20&algo=nmf

NeMOlink12: https://nemoanalytics.org/p?p=p&l=NeocortexDevoHsInVitro&c=MammCtxDev.jNMF.p7&algo=nmf

NeMOlink13: https://nemoanalytics.org/p?p=p&l=NeocortexDevoHsInVitro&c=MammCtxDev.jNMF.p40&algo=nmf

NeMOlink14: https://nemoanalytics.org/p?l=NeocortexDevoHsInVitro&g=EOMES

NeMOlink15: https://nemoanalytics.org/p?p=p&l=NeocortexDevoHsInVitro&c=HsCtxLayer.jNMF.p20&algo=nmf

