## Supplemental Figures for "A Curated Compendium of Transcriptomic Data for the Exploration of Neocortical Development"

**Supplemental Materials**

**Supplemental Table 1:** Gene loadings and enrichment analyses related to the jointNMF() decomposition of scRNAseq data from mouse (La Manno2021) macaque (Micali 2023), and human (Trevino 2021) fetal neocortical tissue detailed in Figures 1 & 2 (p7CtxDev).

**Supplemental Table 2:** Gene loadings and enrichment analyses related to the jointNMF() decomposition of scRNAseq data from mouse (La Manno2021) macaque (Micali 2023), and human (Trevino 2021) fetal neocortical tissue detailed in Figure 3 (p40CtxDev).

**Supplemental Table 3:** Gene loadings and enrichment analyses related to the jointNMF() decomposition of snRNA-seq (SMART-seq) data from adult human neocortical tissue (Jorstad 2023; Bakken 2021) detailed in Figure 4 & 5 (p20CtxLayer).

**Supplemental Figure 1 - related to main Figure 1**

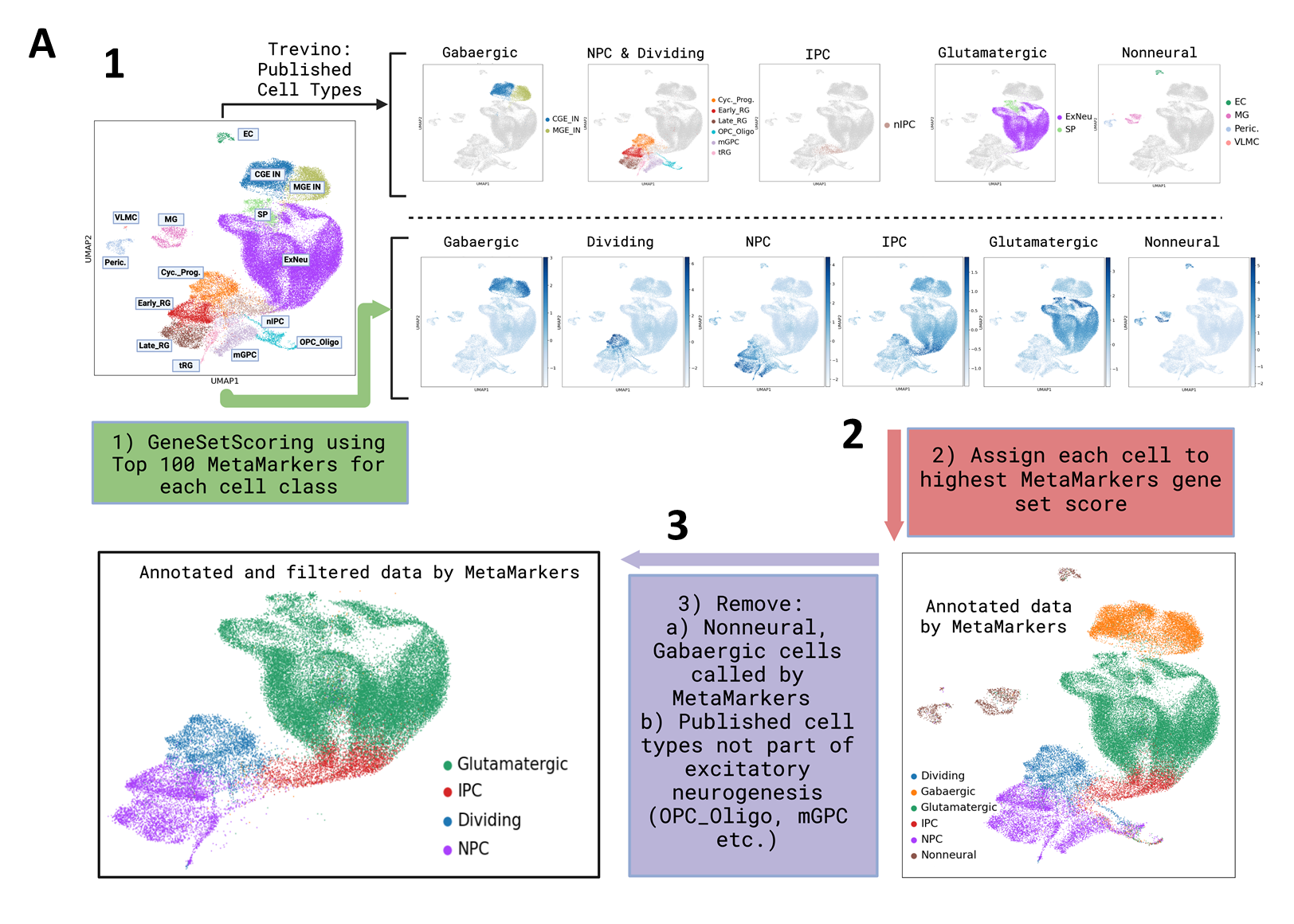

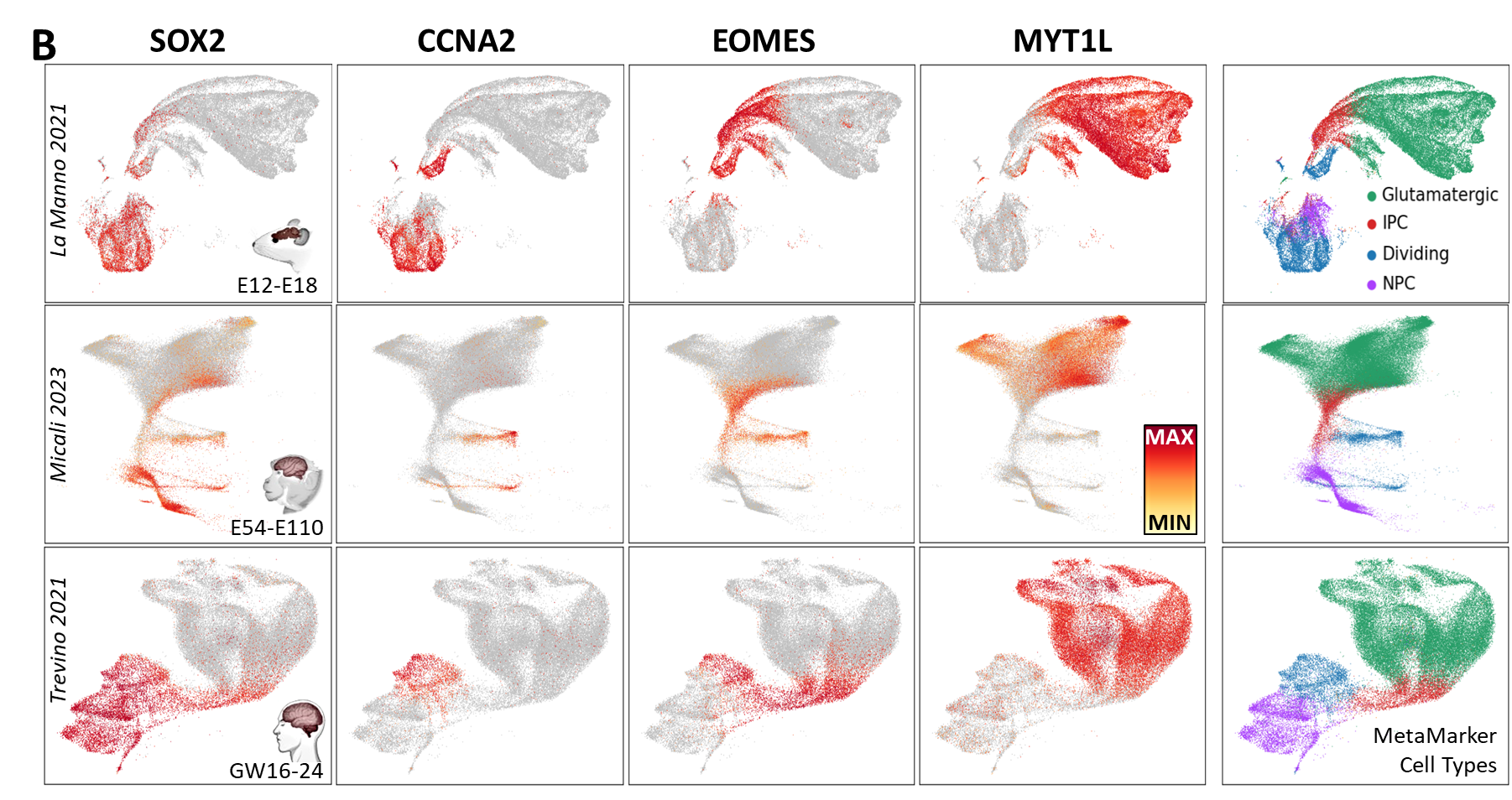

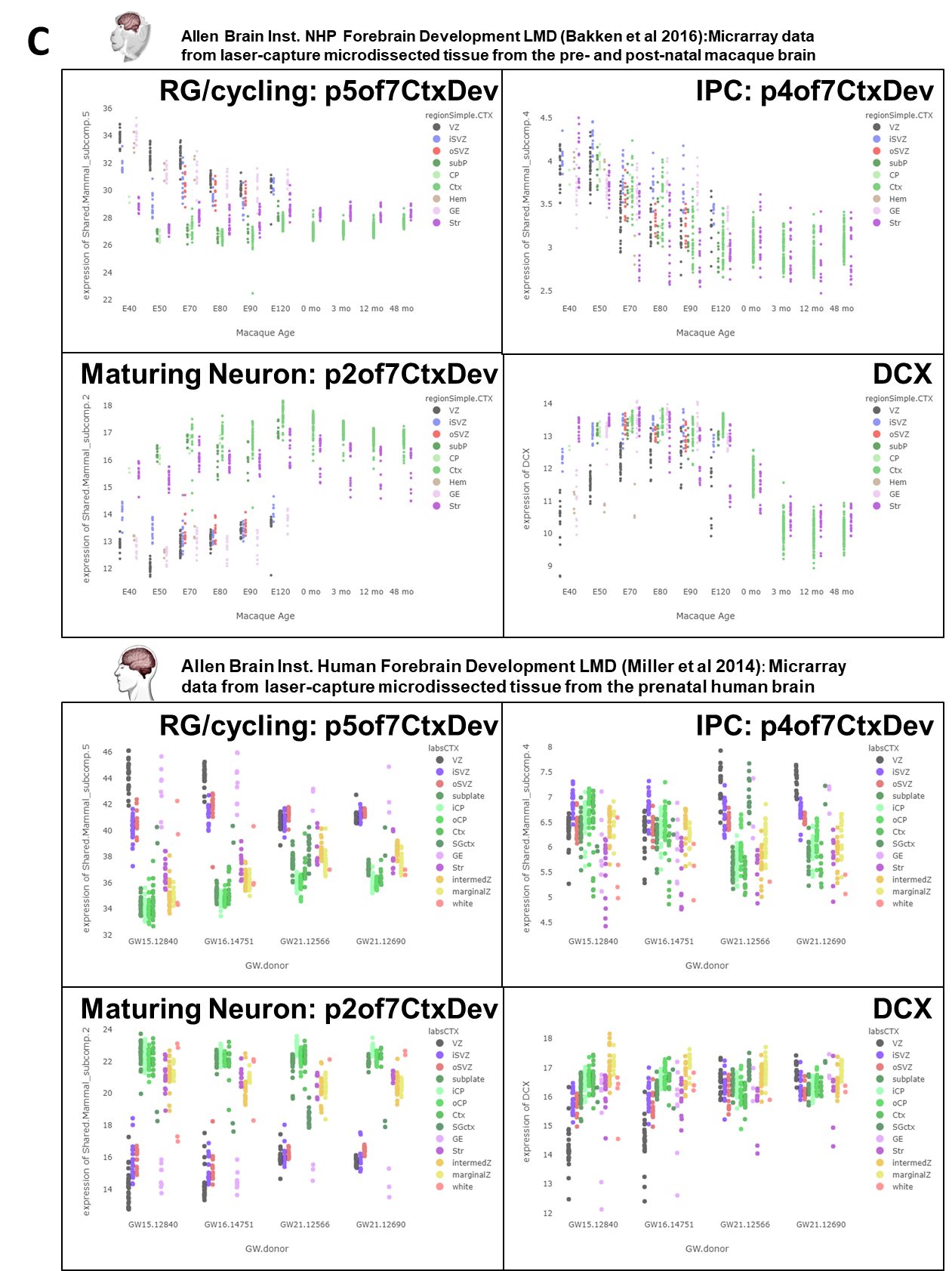

**Supplemental Figure 1 - related to main Figure 1: A]** Example of MetaMarkers classification procedure on Trevino 2021 dataset used for cell type calls in Figure 1 and 3. (1) Large panel and top row: Original author-published UMAP colored by author-curated cell types. Bottom row: Published UMAP colored by each of the 6 MetaMarker gene set scores calculated for each cell using the top 100 marker genes for each MetaMarker class (Methods). (2) Each cell was assigned to the MetaMarker class with the highest gene set score in that cell. (3) Nonneural and Gabaergic cells called by MetaMarkers classification were removed as well as author labeled cell-types not part of the excitatory neurogenic lineage. Further details regarding the procedure are in Methods and Code. **B]** Colorized UMAP plots for each of the three datasets used in Figure 1 showing the expression of 4 individual genes representative of broad cell types in neurogenesis: SOX2 in neural progenitors, CCNA2 in cycling progenitors, EOMES in intermediate progenitors, and MYT1L in new neurons. UMAPS in the right most column are colored by the MetaMarker cell type calls as made in panel A. **C]** In Figure 1D we see that the RG/cycling pattern (p5of7CtxDev) progressively declines in progenitors, while the maturing neuron pattern (p2of7CtxDev) increases in progenitors over time and the IPC pattern (p4of7CtxDev) is reduced over time in neurons. Projection of microarray data from laser capture microdissected (LMD) regions of the developing macaque and human brain into these transcriptome dimensions confirm all of these effects. While these effects could have been related to “ambient” RNA effects from unintentionally lysed cells in the scRNA-seq data, their confirmation in the LMD data demonstrates that this is not the case ([NeMOlinkS01](https://nemoanalytics.org/p?p=p&l=NeocortexEvoDevo&c=MammCtxDev.jNMF.p7&algo=nmf)). Rising expression of DCX in the VZ in these same data is a single gene example of the “neuronalization” of progenitors and can also be viewed online at [NeMOlinkS02](https://nemoanalytics.org/p?l=NeocortexEvoDevo&g=DCX).

**Additional note on gene set enrichments in main Figure 1:**	While the neuronal transcriptome patterns most enriched in synapse associated genes (p7 and p2of7CtxDev) show the highest enrichments for ASD, SCHZ, and BD (Figure 1C, consistent with PMID: 38103876, PMID: 35396580, PMID: 30610197, PMID: 29651456), we must take caution in interpreting these enrichment analyses that have become ubiquitous in efforts to use gene expression data combined with GWAS results in order to infer mechanisms of GWAS variants. The basic premise is sound: regulatory GWAS variants can only function to confer risk for disease in cells where they affect gene expression, so we should look at cell types where genes linked to GWAS variants are expressed. This thinking would for instance indicate that we should look to brain rather than liver for the function of ASD, SCHZ, and BD associated variants because affected genes are expressed in brain and not liver. The reasoning breaks down when comparing multiple tissues or cell types which express the risk-associated genes a different levels. There is no more reason to implicate cells that have higher expression rather than lower. As long as the genes are expressed in a cell or tissue, there is a possible functional role. So focus should be on the number of risk genes expressed in a cell type or tissue rather than the level at which it is expressed, but virtually all conventional enrichment testing assesses expression levels. Additionally, given that these are developmental disorders, a focus on the cells where the genes are first expressed may be more effective than where genes are expressed at the highest levels. Even if levels of risk genes are lower in neural progenitors than neurons, it may be there that risk is likely to play out. This is consistent with recent studies showing that genes previously considered to be solely synaptic in function also have much earlier roles in neural progenitor function (PMID: 35585091, PMID: 37946050). Adding to the notion that earlier events may harbor risk mechanisms, a recent study has indicated that genetic effects on gene expression (eQTLs) are strongest during *in utero* brain development before the formation of most neocortical synapses (PMID: 38781368). Another of our recent reports has examined risk in progenitors in detail (PMID: 38915580).

**Supplemental Figure 2 - related to main Figure 2**

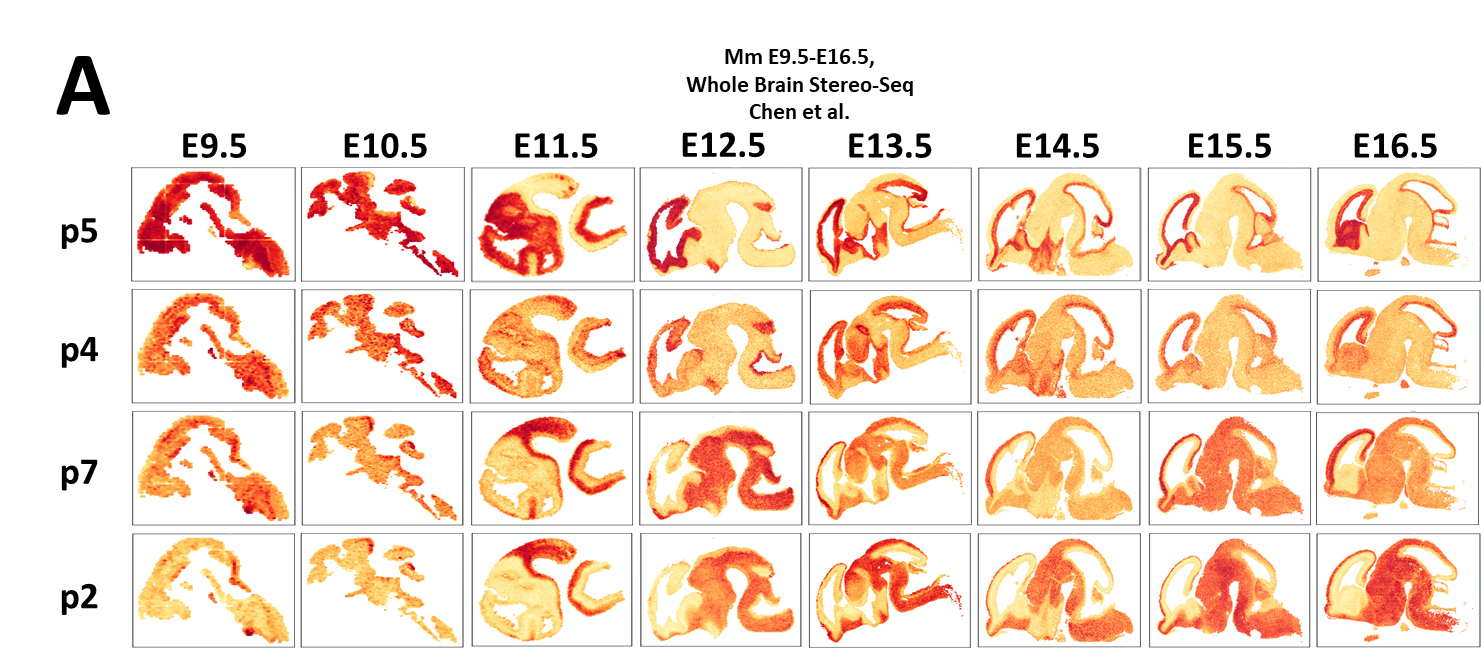

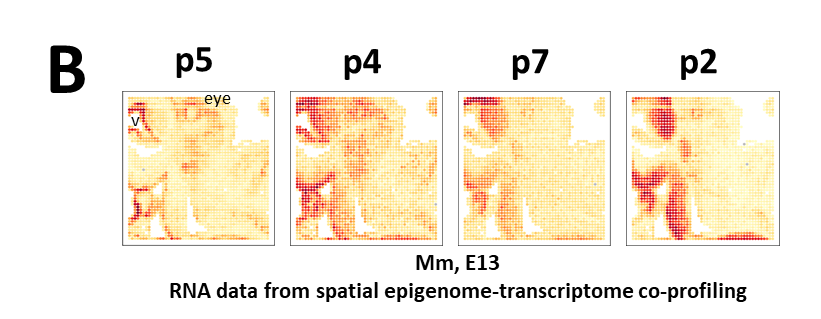

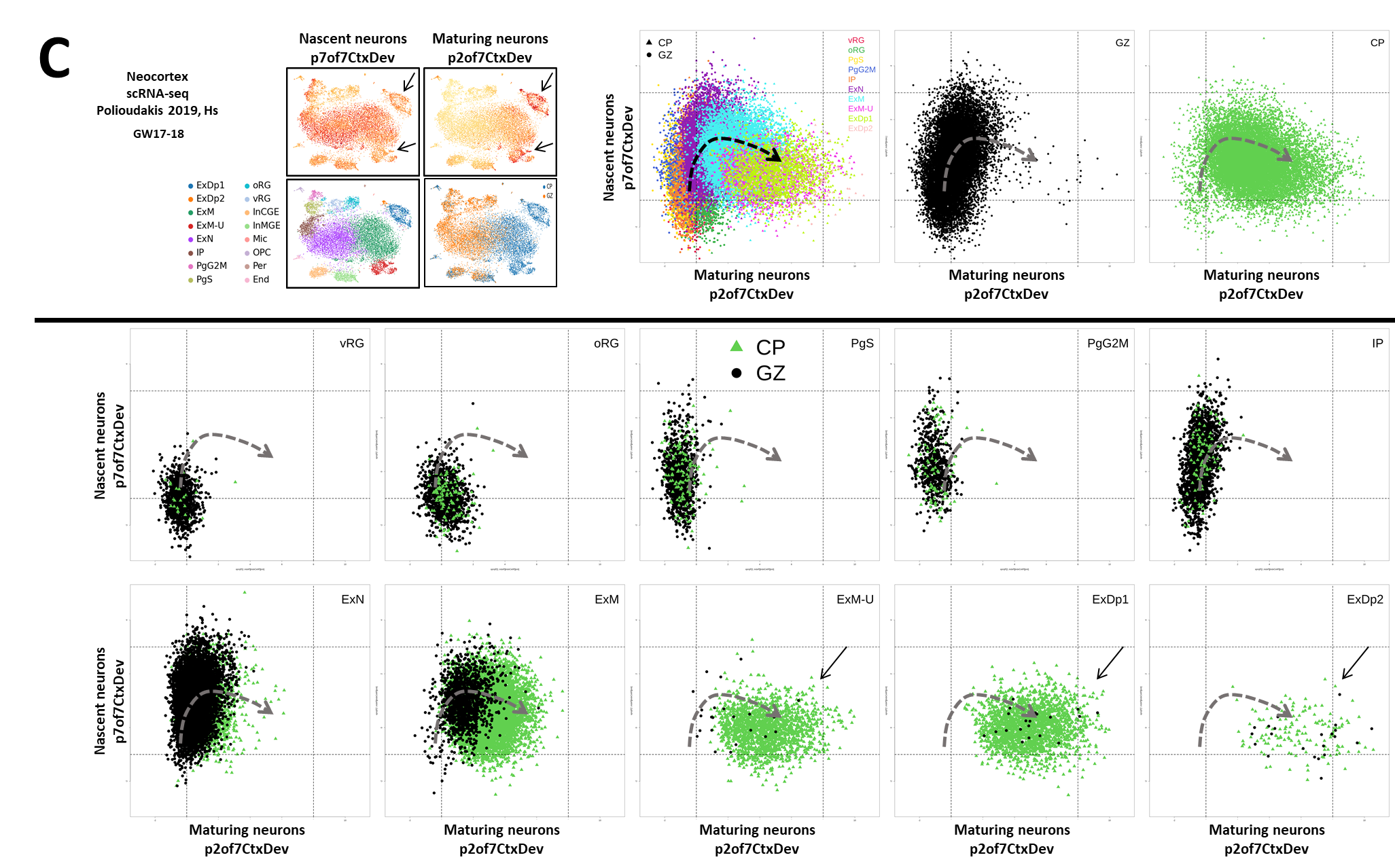

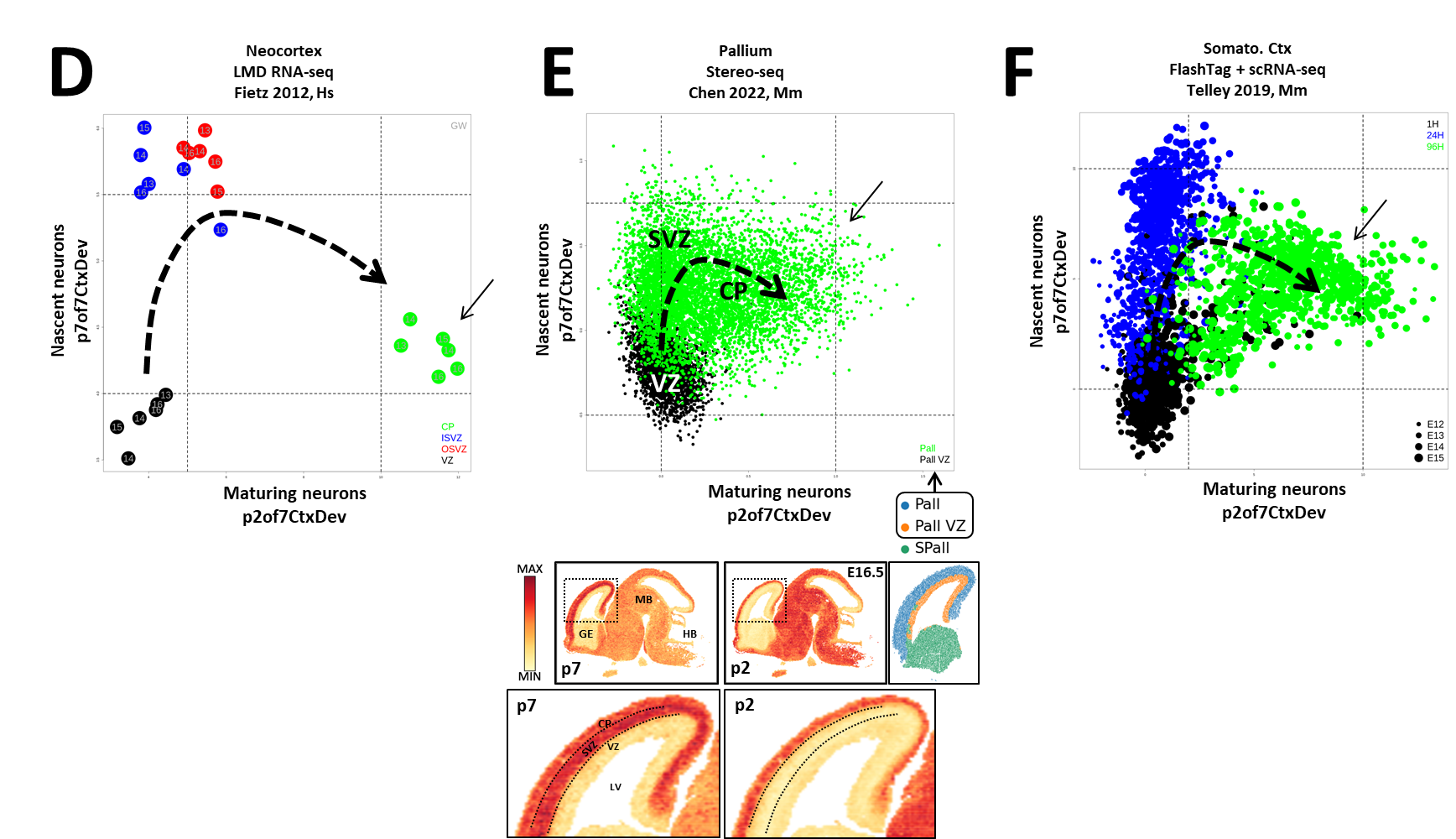

**Supplemental Figure 2 - related to main Figure 2: A]** Visualization of projected embeddings in p5, p4, p7 and p2of7CtxDevo for individual spatial measurements in full sagittal brain slices from E95.-E16.5 in the Chen 2022 dataset (PMID: 35512705). These sections indicate clearly that both the progenitor and neuronal transcriptome signatures are not specific to the neocortex, but rather are pan-brain signatures of these broad cell types. Many spatio-temporal details can be seen across these sections as the developing brain progresses from a tissue dominated by progenitors to a tissue of post-mitotic neurons. **B]** Projected embeddings in p5, p4, p7 and p2of7CtxDevo for individual RNA measurements from spatial epigenome-transcriptome co-profiling (PMID: 36922587). Here the outward radial progression of cells expressing p5 (RG at the ventricle), to p4 (delaminated IPCs), to p7 (nascent neurons) and finally p2 (maturing neurons) can be seen. V=ventricle, lateral, of the forebrain. **C]** Related to data from Polioudakis 2019 (PMID: 31303374) in Figure 2: Upper left panels show the original author published tSNE with author’s cell type calls and plots colored by p7 (proneural nascent neuron signature) and p2of7CtxDev (maturing neuron signature). Original author cell type calls: Ex=excitatory neurons, Dp=deep, N=new/migrating, M=maturing, Ip=intermediate progenitor, Pg=cycling progenitor in S or G2M phase, RG=ventricular (v) or outer (o) radial glia, In=inhibitory neurons of the medial (MGE) or caudal (CGE) ganglionic eminence, Mic=microglia, OPC=oligodendrocyte precursor cell, Per=perictye, End=endothelial cell. Three panels at upper right show projected embeddings of cells in the excitatory neurogenic trajectory (according to author cell type calls) as they move through the transcriptomic dimensions defined by p7 and p2of7CtxDev, first colored by individual cell types and then separately by where the single cells were dissected from in the developing cortex. GZ=germinal zones, CP=cortical plate (both these dissections included portions of the intermediate zone – this is especially relevant to panel G in main Figure 2, where the intermediate zone was dissected separately in the macaque developing neocortex and show the descent pf p7 as p2of7CtxDevo begins to rise). Lower two rows of panels show individual cell types in the p7 v p2 transcriptomic space (colored by their dissection of origin) as they progress through neurogenesis. Throughout panel A, vertical and horizontal dashed lines are in identical positions in each plot to enable the comparison of p7 and p2of7CtxDev pattern values across cell types in different plots. **D]** Scatter plots of projected embeddings in p7 and p2of7CtxDevo for individual samples from LMD coupled RNA-seq in the developing human neocortex from the Fietz 2012 (PMID: 22753484) dataset. LMD=laser microdissection. CP=cortical plate, OSVZ=outer subventricular zone, ISVZ=inner subventricular zone, VZ=ventricular zone. **E]** Scatter plot of projected embeddings in p7 and p2of7CtxDevo for individual spatial measurements in the Chen 2022 dataset (PMID: 35512705). The spatial segmentation of pallium (Pall) and pallium ventricular zone (Pall VZ) used by the original authors is used here to color these individual spatial measures in the scatterplot. These same projected embeddings are also shown in NeMO across the spatial section beneath the scatter plot. **F]** Scatter plot of projected embeddings in p7 and p2of7CtxDevo for Flash-Tag labeled (birth-dated) single-cells in the Telley 2019 dataset (PMID: 31073041). The time from terminal division is represented by color in the scatter plot. In panels **C-F**, arrows indicate maturing neurons, where p7of7CtxDev has descended and p2 is highest. Along with main Figure 1G, this temporal and spatial understanding of new-born neurons as they move through these transcriptomic dimensions is central for their use in Figures 6 and 8 where we harness these same signatures to explore the protracted maturational trajectories of adult neuronal signatures.

**Supplemental Figure 3 - related to main Figure 3**

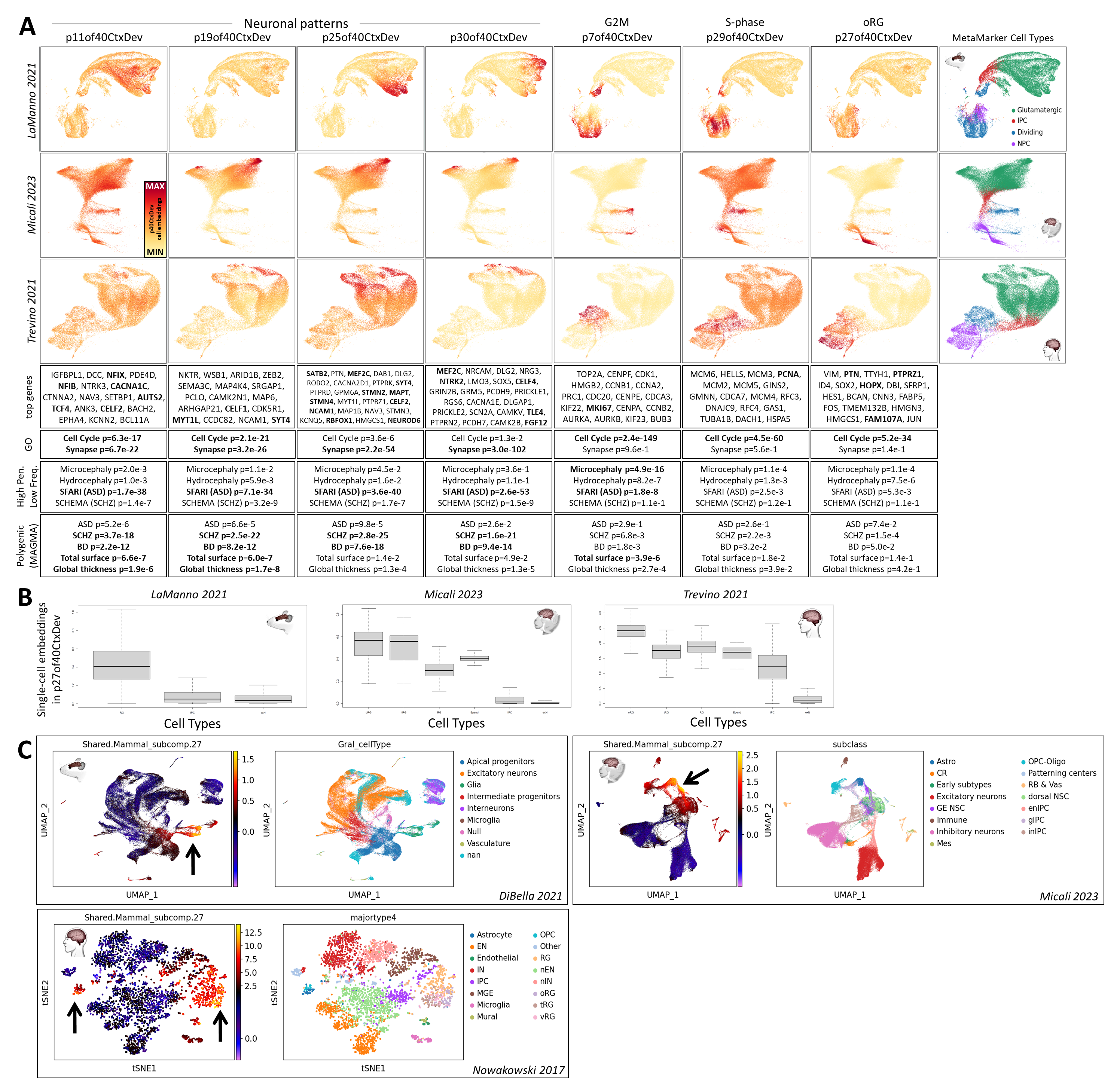

**Supplemental Figure 3 - related to main Figure 3: A]** Single-cell embedding UMAP plots of all three mammalian neocortical datasets for 7 selected patterns (4 patterns that appear in differing subsets of neurons and 3 that define transcriptomic programs within progenitors) from the full set of 40 (p40CtxDev) along with individual genes and gene set enrichments for each. Many details can be seen in this higher resolution decomposition of the mammalian neocortical neurogenesis datasets. Somewhat paradoxically, gene loadings for neuronal patterns p11of40CtxDev and p19of40CtxDev show strong enrichments for both synaptic genes and genes involved in the cell cycle (Figure S3A). This again highlights the fact that transcriptomic programs span cell populations that are often defined as discrete non-overlapping classes, but which in fact share many molecular attributes. Interestingly, in addition to highly significant enrichments for neuropsychiatric diseases, enrichments in the genetics of brain structure are greatest in these patterns that appear to bridge proliferative and post-mitotic states in neurogenesis. Patterns p25of40CtxDev and p30of40CtxDev have much less cell cycle involvement and show even stronger enrichments in synaptic as well as disease associated gene sets (Figure S3A). Individual genes with especially high loadings in these patterns include well-known neuropsychiatric risk genes, including RBFOX1 and MEF2C, many synaptic genes, and layer-defining transcription factors (TFs), including SATB2 and TLE4. Hence, neuronal patterns defined in this higher resolution decomposition show enrichment in genes linked to human brain structure and disease, with hits of similar significance to those found in the low resolution decomposition (Figure 1), but here with greater precision in defining the subpopulations of genes and cells that are involved with these associations. Also in this higher resolution decomposition, transcriptomic elements of neural progenitors and the cell cycle have been separated in greater detail. Two new patterns which span distinct subpopulations of both RGs and IPCs are shown in Figure S3. Both the ranking of individual marker genes among the patterns’ gene loadings (G2M marker MKI67 ranked #22 in p7of40CtxDev and S-phase marker PCNA ranked #5 in p29of40CtxDev) and enrichment analysis (G2M checkpoint in p7of40CtxDev, p=4.9e-76; DNA replication in p29of40CtxDev, p=2.1e-19) clearly indicate that these patterns are present specifically in the transcriptomes of cells in G2M (p7of40CtxDev) and S-phase (p29of40CtxDev) of the cell cycle. In the previous 7 pattern analysis (p7CtxDev), the RG and cycling cell pattern p5of7CtxDev showed enrichment for genes involved in monogenic microcephaly (p=4.1e-9), but only a modestly significant enrichment for genome-wide association with total cortical surface area (p=0.001). The G2M pattern p7of40CtxDev is present in a much more limited subpopulation of progenitors than p5of7CtxDev, yet showed an even greater enrichment for both monogenic microcephaly (p=4.9e-16) and total cortical surface area (p=3.8e-6). In the Micali 2023 macaque data this dissection of the cell cycle can be seen independently in radial glia (lower horizontal loop with distinct areas of high p7 and high p29) and intermediate progenitors (upper horizontal loop with distinct areas of high p7 and high p29). These results indicate that by elevating the resolution of transcriptome decomposition, it is possible to identify increasingly specific transcriptomic elements within cell subtypes and their association with brain structure and disease. Additionally, gene loadings for this oRG signature were enriched in genes involved in cholesterol homeostasis (p=6.2e-12). Along with the canonical oRG markers, KLF6, a gene recently implicated in evolutionarily new mechanisms in cholesterol metabolism specifically in human oRG cells (doi.org/10.1101/2023.06.23.546307) is also among the top ranked genes in p27of40CtxDev. Supplementary Table 3 contains the full set of gene loadings across all 40 patterns and Supplementary Table 4 contains the full set enrichments across all 40 patterns. **B]** Boxplots of projected single-cell embeddings for p27of40CtxDev (oRG) separated by cell type calls made by original authors. For LaManno 2021 (PMID: 34321664) and Micali 2023 (PMID: 37824652) the original author cell type calls were used. Trevino 2021 (PMID: 34390642) did not make detailed cell type calls (including no oRG calls). Therefore, we added some detail to these calls for both this boxplot and the CellOracle analysis in the main Figure 3 (See “Trevino et al 2021 Cell type reannotation and CellOracle analysis” section of Methods). **C]** p27 has high expression levels in gliogenic precursors and astroglia in mouse (PMID: 34163074), macaque (PMID: 37824652), and human (PMID: 29217575) cortex. This is consistent with our recent observations employing orthogonal methods, which indicate that the first evolutionarily components of the oRG transcriptomic program arose in gliogenic precursors of the rodent-primate ancestor before expanding and driving the evolution of oRG cells in the primate lineage (PMID: 37383947). Explore more at [NeMOlinkS03](https://nemoanalytics.org/p?p=p&l=NeocortexEvoDevo&c=MammCtxDev.jNMF.p40&algo=nmf) by selecting p27. Plots in panels A and C are screen captures from NeMO Analytics.

**Supplemental Figure 4 - related to main Figure 4**

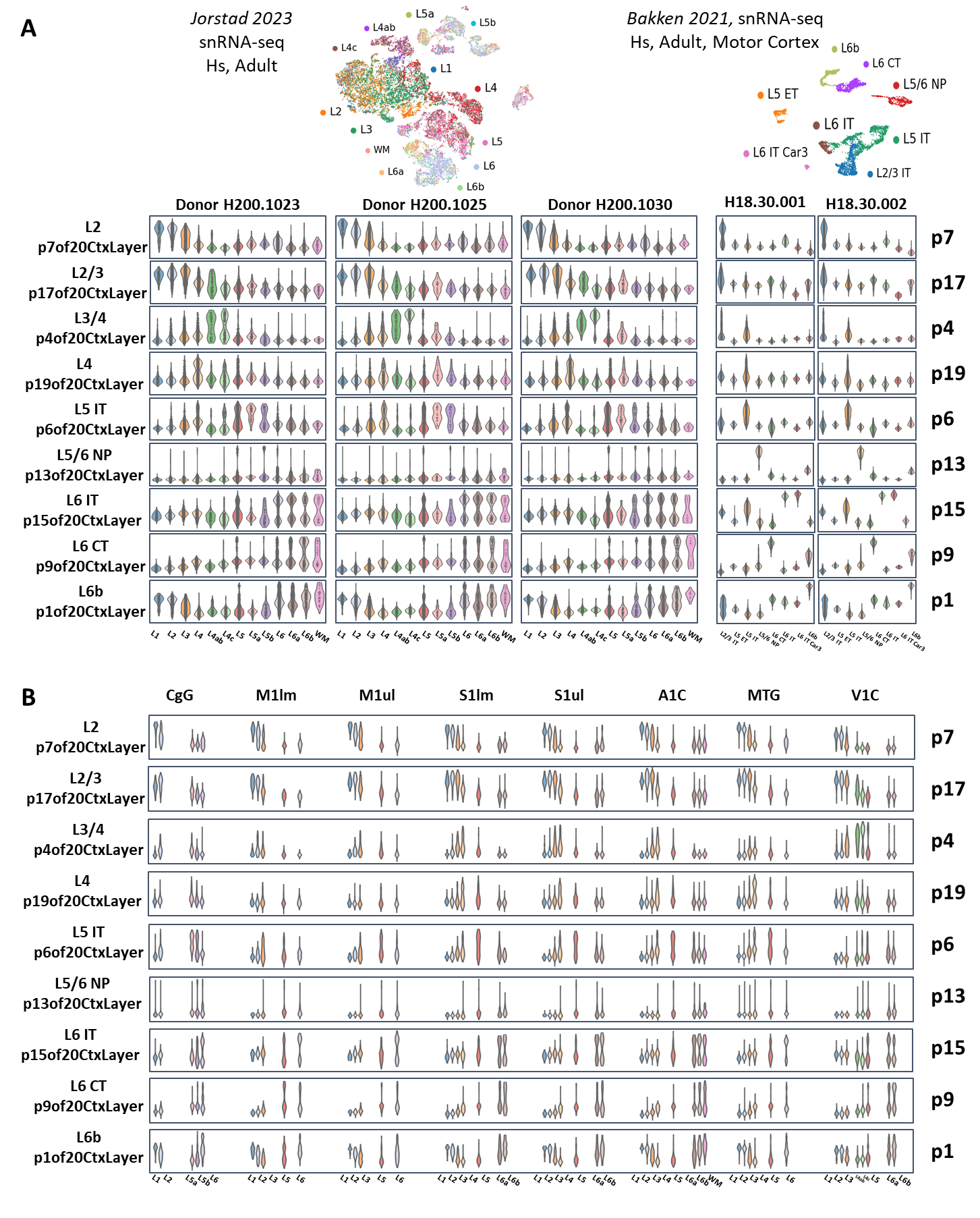

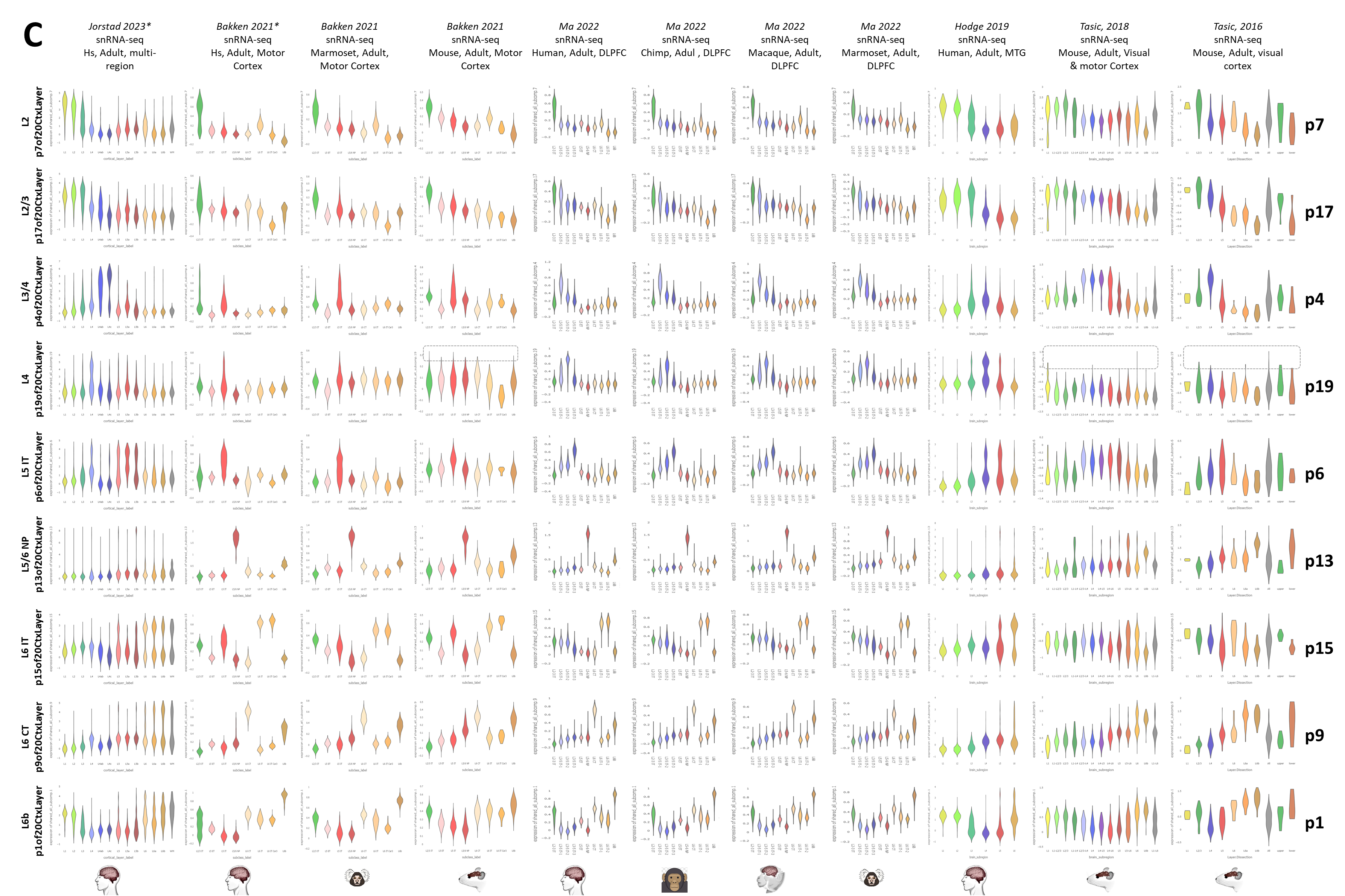

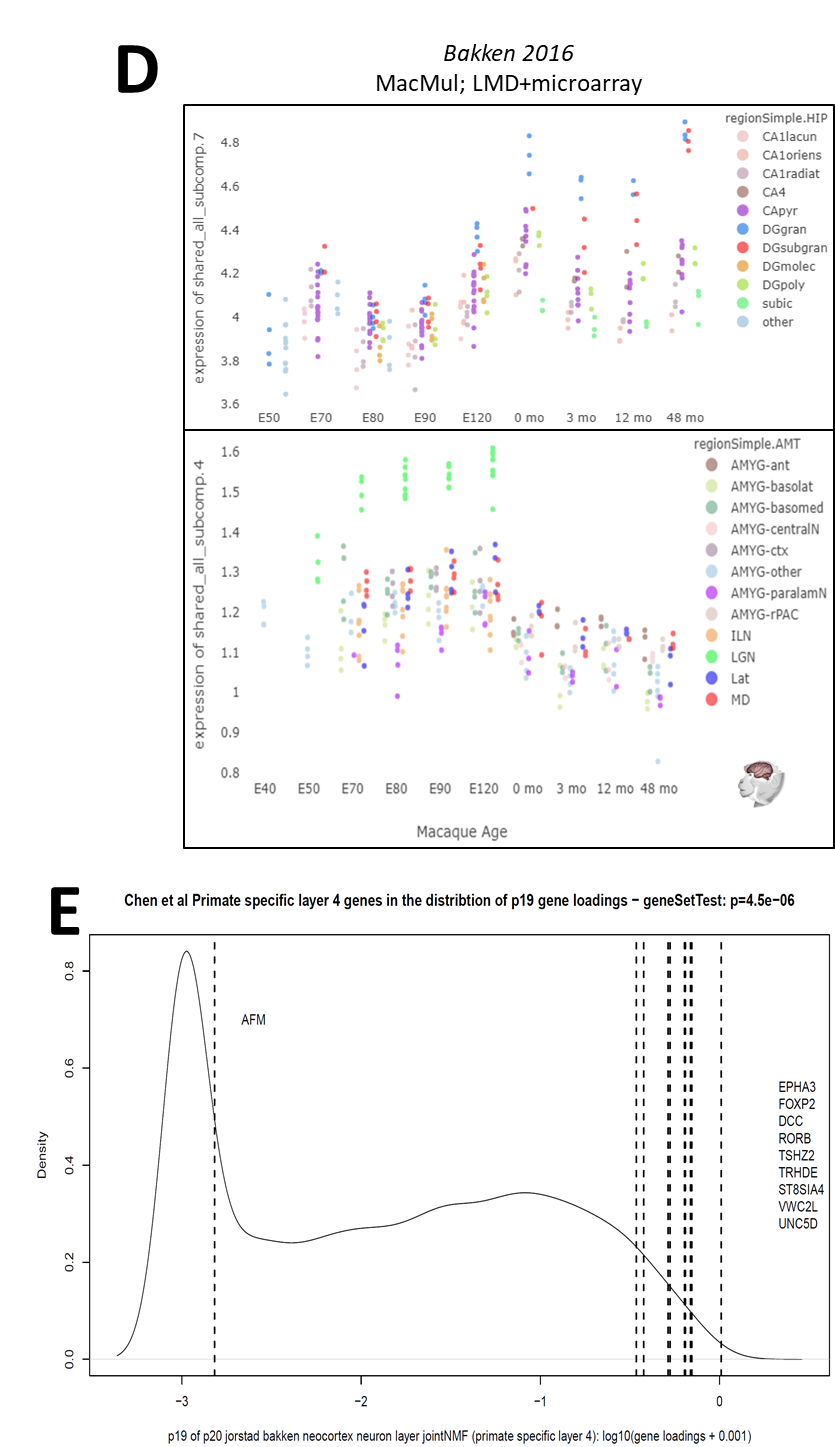

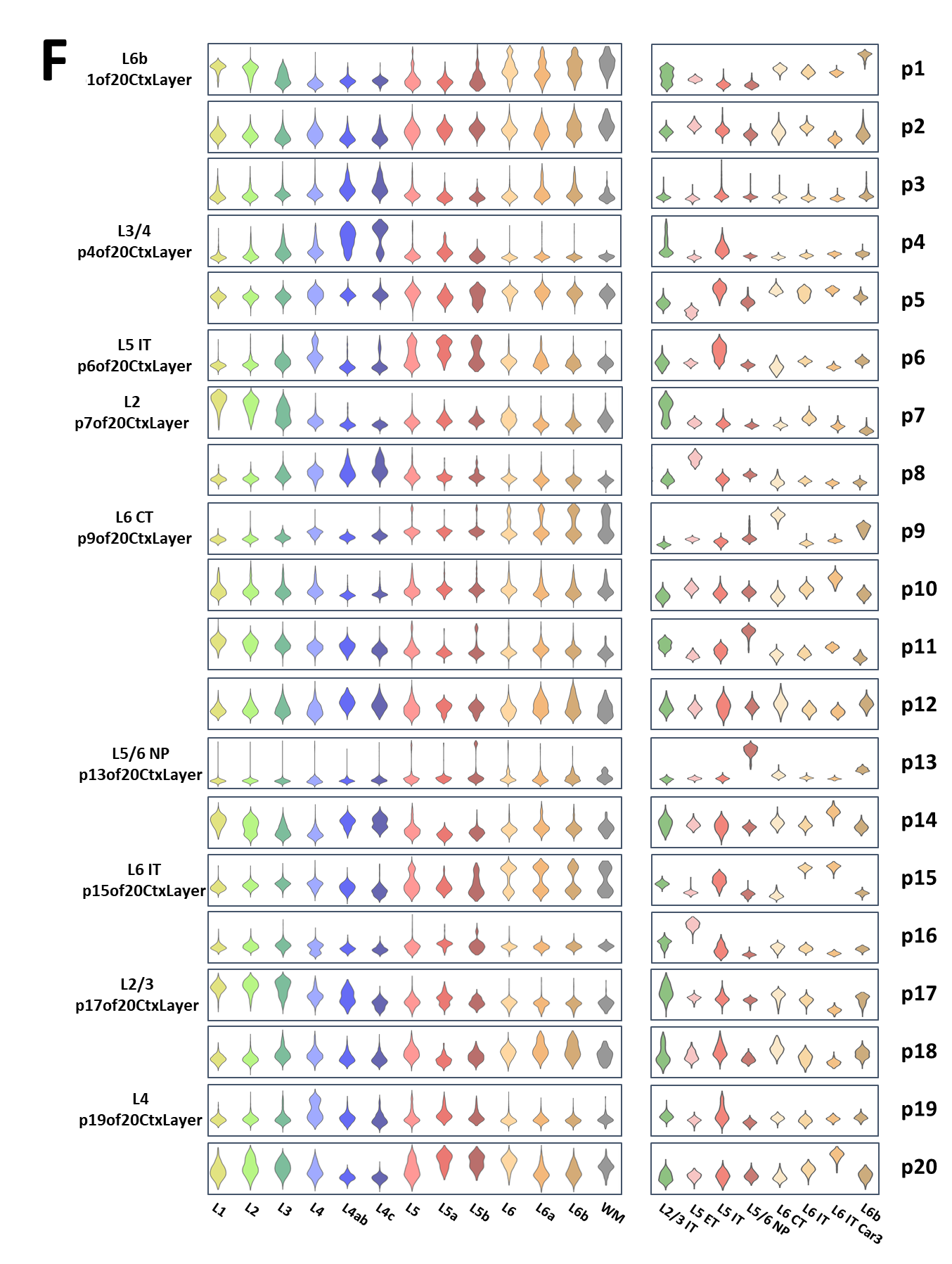

**Supplemental Figure 4 - related to main Figure 4: A]** Violin plots of cell embeddings for the two adult human snRNA-seq (SMART-seq) datasets used in the jointNMF() decomposition in Figure 4 (Jorstad 2023, PMID: 37824655 & Bakken 2021, PMID: 34616062). Embeddings for 9 of the 20 total transcriptomic signatures (p20Layers) are shown. These are the same 9 focused on in main Figures 4 &5, where projections in other datasets are shown. Here embeddings from individual donors are shown separately to show consistency across samples. **B]** Violin plots of cell embeddings from the same Jorstad 2023 (PMID: 37824655) data as in panel A, here rearranged to show embeddings from individual regions to show consistency across neocortical areas. **C]** Violin plots for a wide collection of snRNA-seq datasets demonstrating the species and laminar specificity of the transcriptomic signatures from adult human cortex focused on in Figure 4 & 5. Dashed boxes indicate where the primate and human specific layer 4 pattern, p19of20CtxLayer, is absent in the mouse neocortex. Asterisks (*) indicate the two datasets used in the original decomposition. **D]** Projection of LMD coupled microarray expression data from non-cortical regions of the developing macaque brain into p7 and p4of20CtxLayer which showed shared signatures with hippocampal and thalamic neurons, respectively, in mouse spatial transcriptomic data in main Figure 4. The LMD data confirm that these transcriptomic signatures are also shared across these same brain regions in macaque, with p7 at high levels specifically in the dentate gyrus granule layer, and p4 high in the lateral geniculate nucleus of the thalamus. These projections clearly demonstrate that multiple human adult laminar neuron transcriptome patterns in the neocortex are shared with neurons in diverse regions of the brain and these shared programs are conserved across mammals, consistent with recent scRNA-seq mapping of neuronal types across the cortex and hippocampus in the mouse brain (PMID: 34004146) where numerous neuronal types were observed to be common across these two brain structures. **E]** Density plot of gene loadings in the p19of20CtxLayer transcriptomic signature of primate and human specific layer 4 neurons. The loadings in for the 11 genes reported by Chen 2023 (PMID: 37442136) to be specifically in layer 4 neurons of primates and humans are shown as vertical dashed lines and show significant enrichment among high p19 loadings in a Wilcoxon rank sum test (implemented via the geneSetTest() function in the limma package in the R statistical language; p=4.5e-6). **F]** The complete set of 20 transcriptomic patterns in the p20CtxLayer decomposition, shown as cell embeddings aggregated across donors and regions in the Jorstad 2023 (PMID: 37824655) and Bakken 2021 (PMID: 34616062) data that were used in the decomposition. Original author cell type calls were used.

**Supplemental Figure 5 - related to main Figure 5**

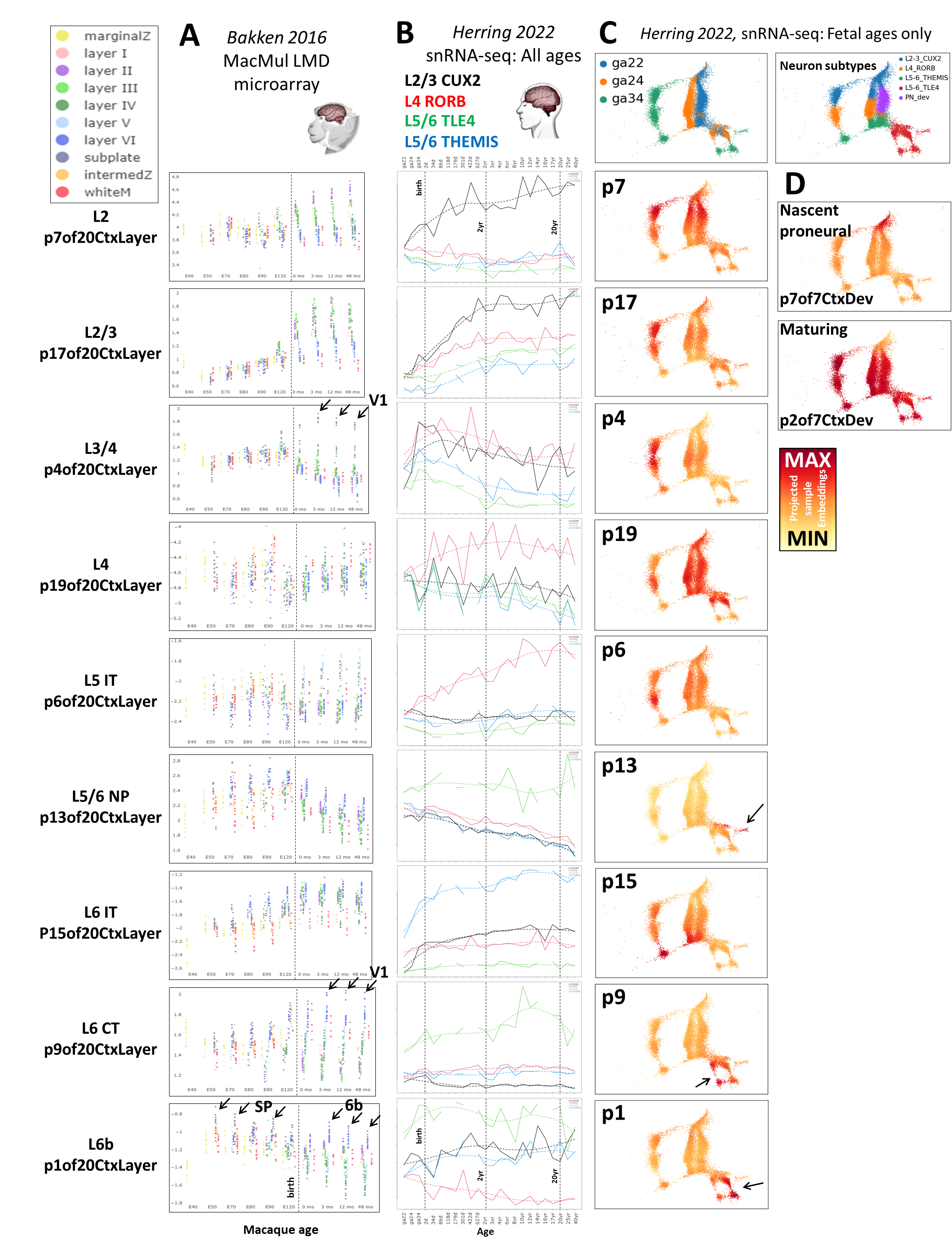

**Supplemental Figure 5 - related to main Figure 5: A]** Projection of macaque LMD-coupled microarray data into the p20CtxLayer patterns. The laminar specificity and protracted timescales of these maturational trajectories in the non-human primate are made clear in the laser capture microdissection (LMD)-coupled expression analysis spanning pre- and post-natal macaque neocortical development from Bakken 2016 (PMID: 27409810). Additional observations in this unique dataset include the linking of fetal subplate and mature layer 6b molecular identities in p1of20CtxLayer (consistent with PMID: 22628460), V1-specific enrichment of the p9 layer 6, and p4 layer 3/4 identities, all of which play out over years of postnatal development (arrows). These transcriptomic signatures can also be explored online (including individual sample annotation in the LMD data) at: [NeMOlinkS04](https://nemoanalytics.org/p?p=p&l=AdultNeoctxLayers&c=HsCtxLayer.jNMF.p20&algo=nmf). **B]** Line plots of the mean projected single-cell embeddings in neurons across pre- and post-natal life for each adult layer-specific neuronal pattern (9 of 20 patterns in p20CtxLayer) in the Herring 2022 (PMID: 36318921) snRNA-seq data split into 4 neuron subtypes identified by the original authors. Solid lines join means at each age/donor. Dashed lines are local fits to the individual age means made with the loess() function in the R statistical language. **C]** Originally published UMAP of fetal neurons colored by projected single-cell embeddings for each adult layer-specific neuronal pattern (9 of 20 patterns in p20CtxLayer) in the Herring 2022 (PMID: 36318921) snRNA-seq data. The adult laminar-specific neuronal transcriptomic signatures (especially the deep-layer signatures) have strikingly clear specificity in particular cell populations at these early stages of development. For example, the 3 patterns which show high levels in TLE4+ neurons (p1, p9, and p13of20CtxLayer), all show high levels in clearly distinct sub-populations of the neurons classified as TLE4+ by Herring 2022 (arrows). While following the general pattern of building laminar-specific expression over time within a particular neuronal type, two of the three patterns in TLE4+ neurons, p1of20CtxLayer (L6b) & p13of20CtxLayer (L5/6 NP), display particularly unique maturational trajectories (see also main Figure 5B). In fetal and early postnatal development, these patterns each rise specifically within distinct sub-populations of TLE4+ neurons and fall in other TLE4+ neurons. Interestingly, levels of p13 fall across development and throughout adulthood in all neuron types with the exception of the specific TLE4+ L5 NP subtype in which they are finally enriched (se also main Figure 5B). Hence, while all layer identities arise from increasing expression levels in a specific neuronal cell type, some also emerge via repression in other neuronal types. **D]** The Herring 2022 (PMID: 36318921) dataset captured very few nascent neurons (high in p7of7CtxDev & low in p2of7CtxDev) and many maturing neurons (low in p7 and high in p2). This is perhaps due to tissue dissection bias or other factors that we have not determined. Comparing panel C to plots in panel D, it appears that the adult layer-specific neuronal transcriptomic signatures begin to emerge in fetal neurons that have shut down p7 and induced p2, as earlier analysis of patterns p7of7CtxDev & p2of7CtxDev in Figure 2 & S2 indicated. This is explored in much greater detail in Figure 6 & S6. With the exception of the line plots and additional labels, this entire figure was created from NeMO Analytics screen captures. Original author cell type calls were used.

**Supplemental Figure 6 - related to main Figure 6**

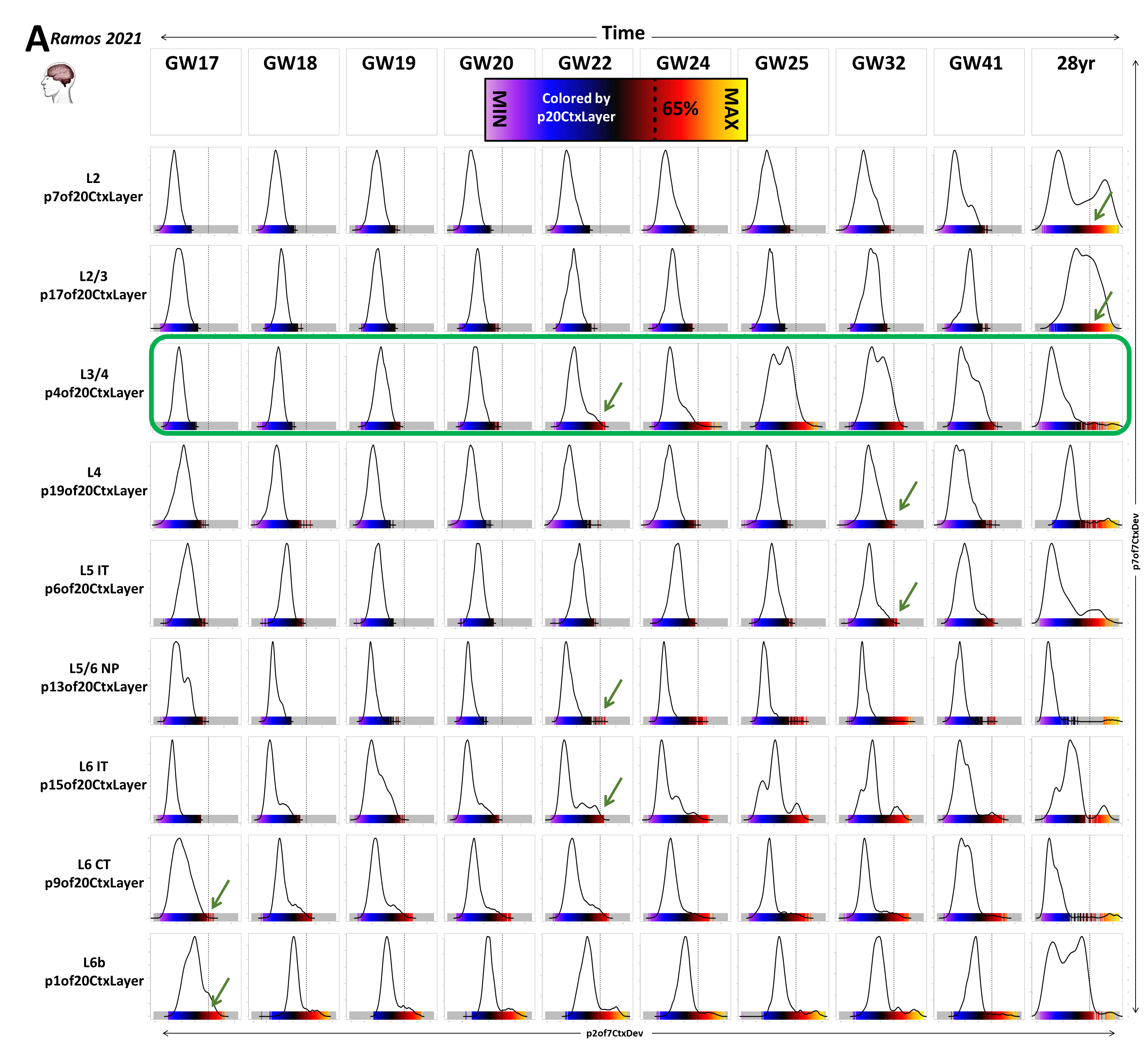

**Supplemental Figure 6 - related to main Figure 6: A]** This collection of plots is constructed just as the matrix of plots in Figure 6, with each mature neuronal pattern in a row and developmental ages across the columns, but here focusing on only the emergence of the adult transcriptomic signatures. The density plots show the distribution of projected single-cell embeddings for each of the adult neuronal transcriptional signatures at each age in fetal development and a single adult time point in the Ramos 2021 (PMID: 36509746) data. Each individual single-cell embedding is plotted as a vertical color strip beneath the density plot. Green arrows indicate the age at which cells first begin to surpass 65% of the maximal level for each signature. In the color scale here (same as in Figure 6) this is where the coloring of samples begins to turn visibly red. As observed in Figure 6, the conserved layer 4 pattern, p4of20CtxLayer, appears earlier than other cells at similar laminar positions (green box). As in Figure 6, this includes only cells of the excitatory neurogenic lineage. To remove low quality cells, in both Figure 6 and here in Figure S6A, we performed an additional QC step, removing cells with fewer than 1250 gene detected or greater than 5% mitochondrial reads.

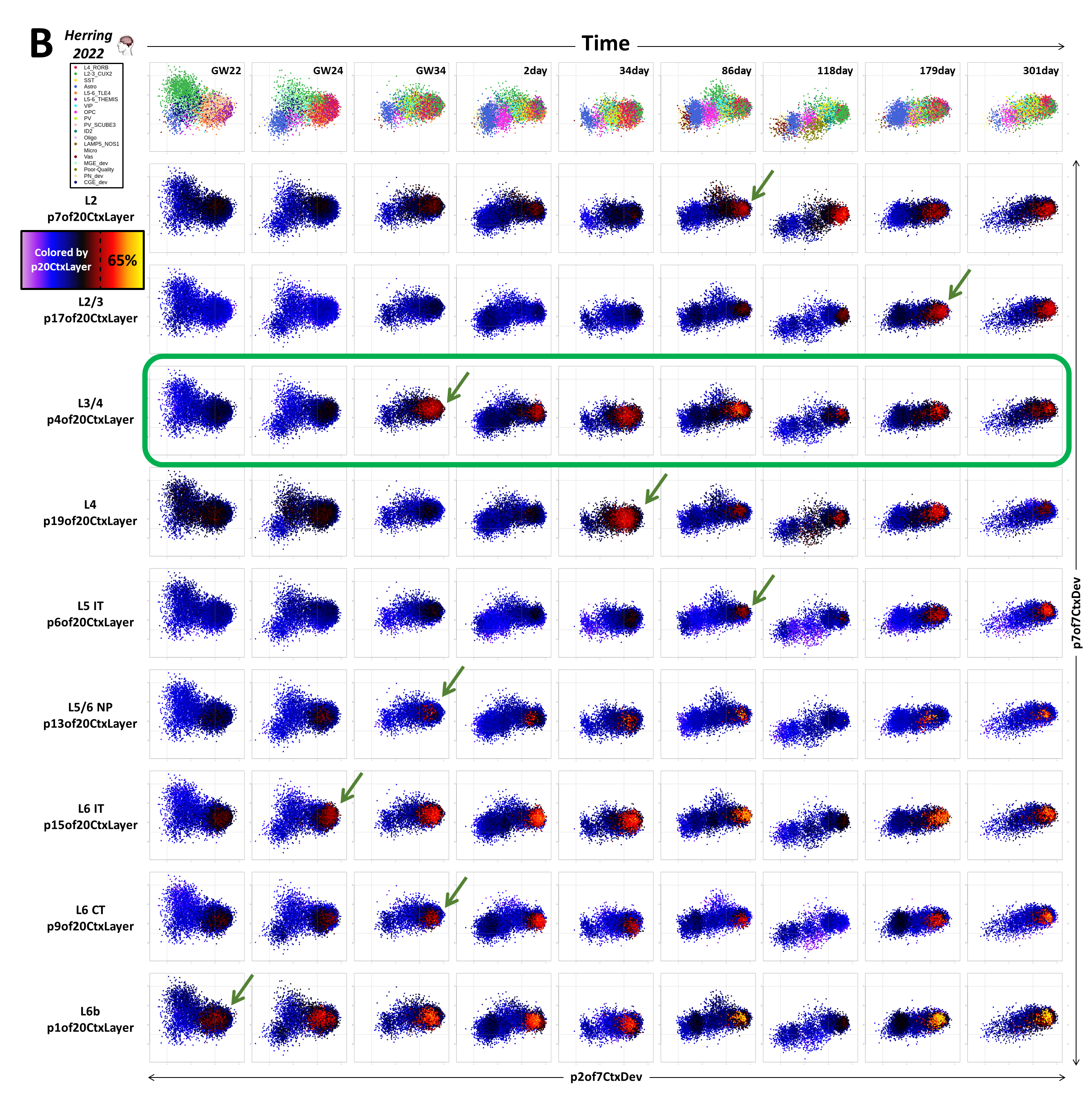

**Supplemental Figure 6 - related to main Figure 6: B]** Recreation of the same plots of projected p7 vs. p2of7CtxDev embeddings colored by projected p20CtxLayer embeddings in Figure 6 using data from Ramos 2021 (PMID: 36509746), here using data from Herring 2022 (PMID: 36318921). Because Herring 2022 (PMID: 36318921) did not make any progenitor cell type calls, we included all cells to ensure we did not miss any of the excitatory neurogenic trajectory. As in the Ramos 2021 (PMID: 36509746) data in Figure 6, here the conserved layer 4 pattern, p4of20CtxLayer, appears earlier than other cells at similar laminar positions (green box). The same 65% threshold that was applied in the Ramos data in Figures 6 and S6A is applied here in the Herring data, however, the Ramos data included many more early fetal time points. This difference in age ranges covered likely inflates the estimated of ages of signature emergence in the Herring data. Original author cell type calls were used.

**Supplemental Figure 7 - related to main Figure 7**

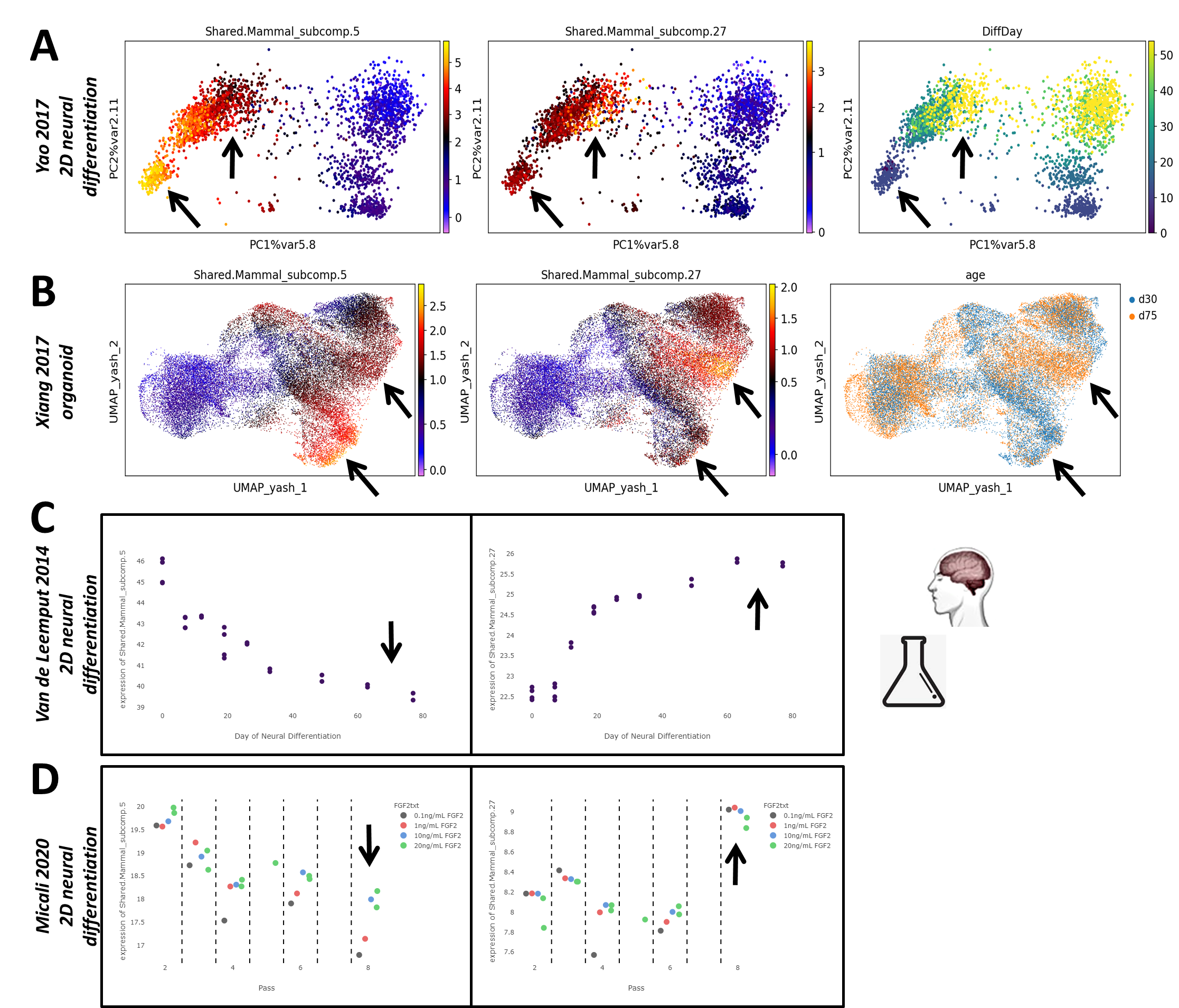

**Supplemental Figure 7 - related to main Figure 7: p5of7CtxDev and p27of40CtxDev differ systematically in their temporal emergence *in vitro*.** As noted using *in vivo* data in Figures 1, 2, and S3, p5of7CtxDev is expressed at particularly high levels in early dividing RG, while p27of40CtxDev is a transcriptomic signature of later oRG and also high in gliogenic precursors that emerge following neurogenesis. Here we show with projection of data from iPSC-derived neural differentiation systems, that these two transcriptional programs emerge in a similar succession *in vitro*: **A]** Yao 2017 (PMID: 28094016) shows p5of7CtxDev highest in neural precursor cells growing in 2-dimensional culture at day 7, and p27of40CtxDev is highest in a small population of precursor cells at day 54. **B]** Xiang 2017 (PMID: 28757360) shows high expression of p5of7CtxDev in precursors at day 35 of organoid culture, while p27of40CtxDev is highest in a distinct population of precursors, later at day 75. **C]** Van de Leemput 2014 (PMID: 24991954) using bulk RNA-seq showed a similar temporal trend, with p5of7CtxDev high early and p27of40CtxDevo high later. **D]** Micali 2020 (PMID: 32375049) using bulk RNA-seq across the passaging of human neural cells shows highest levels of p5of7CtxDev in early passages and highest p27of40CtxDev at later passages. Arrows indicate time points at which the expression of p5of7CtxDev and p27of40CtxDev differ. All panels are screen captures from NeMO Analytics.

**Supplemental Figure 8 - related to main Figure 8**

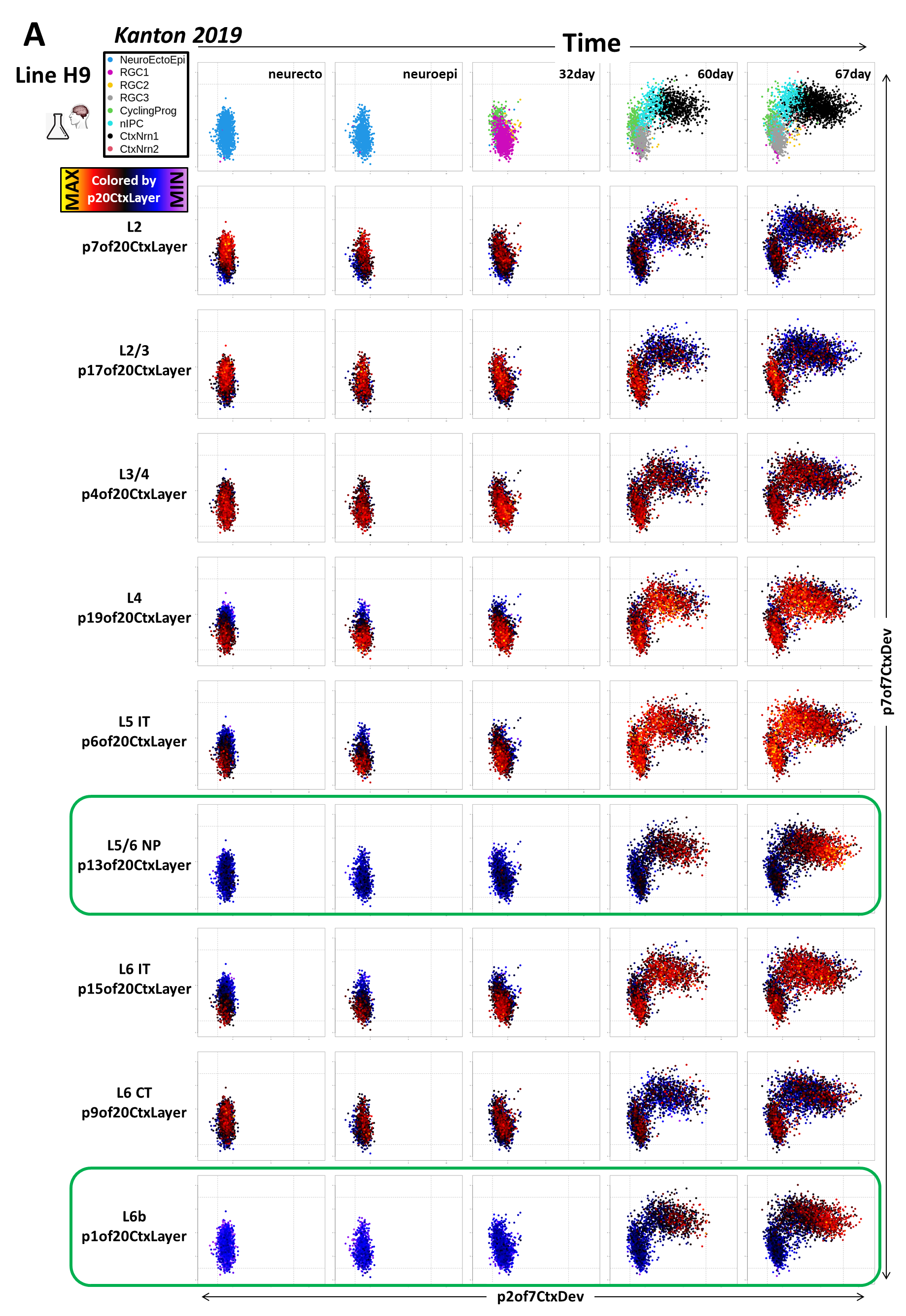

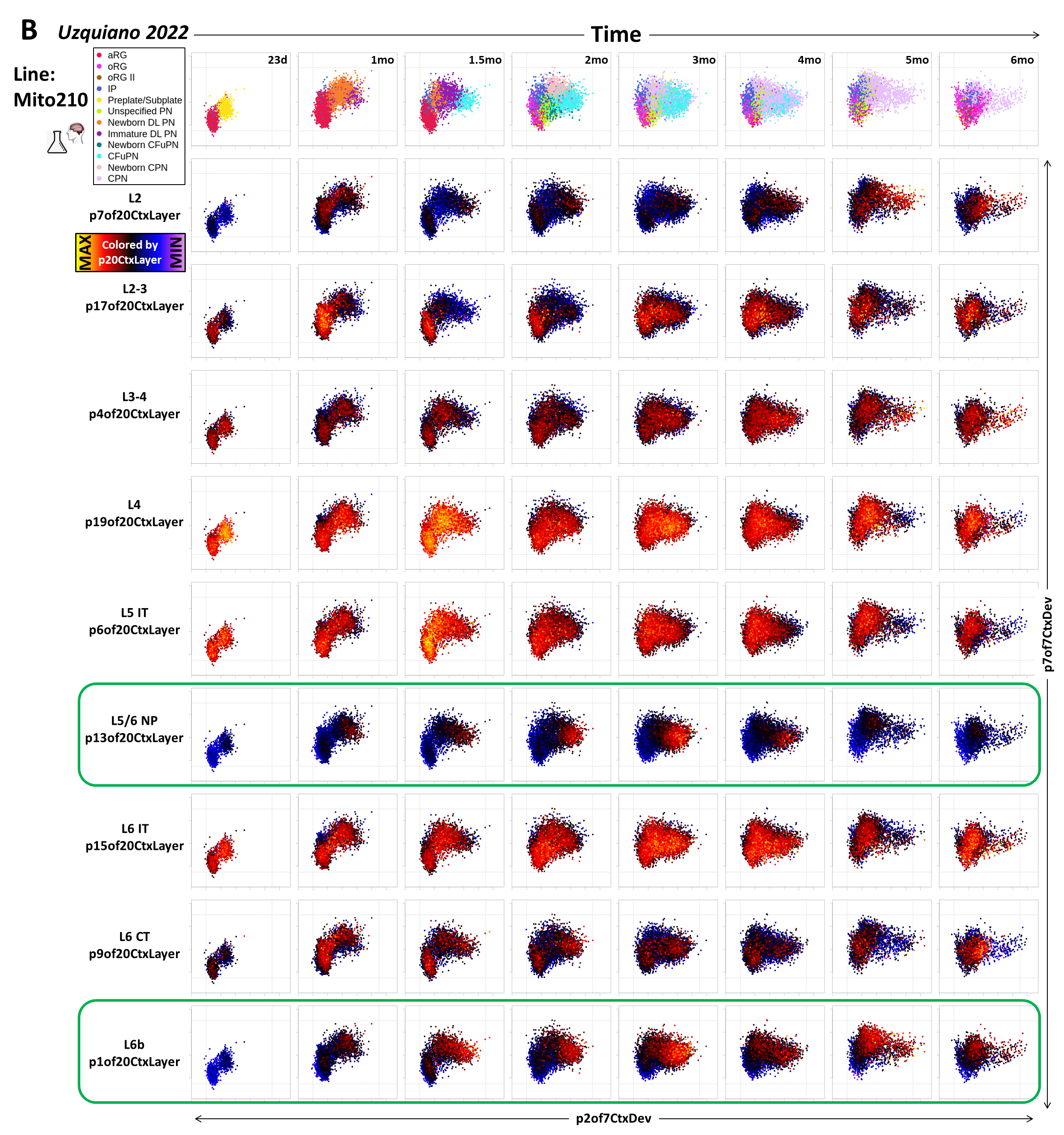

**Supplemental Figure 8 - related to main Figure 8: A]** Main Figure 8 shows the organoid time course for 1 human iPSC line (409b2) from the Kanton 2019 study (PMID: 31619793), here we show the time course for the H9 human ESC line. The same two deep layer transcriptomic signatures appear systematically in the H9 line here as appeared in the 409b2 line (green boxes). **B]** As in both hPSC lines from the Kanton 2019 data, the same two deep layer transcriptomic signatures appear systematically here in the Uzquiano 2022 (PMID: 36179669) organoid time course (green boxes). The Uzquiano 2022 time course is much longer than that in Kanton 2019. The L5/6 NP pattern (p13of20CtxLayer) appears to peak at 3 months, decreasing thereafter and almost disappearing by 6 months. This is likely due to the rise of the gliogenic lineages along with possible necrosis of portions of the organoid that were dedicated to specific neuronal populations. Notably, the L6b pattern (p1of20CtxLayer) also appears to peak at 3 months, but shows a less dramatic decrease thereafter. Original author cell type calls were used: NeuroEctoEpi=neurectodermal and neuroepithelial states, RGC=radial glial cells, CyclingPrg=cycling neural progenitors, nIPC=neuronal intermediate progenitor cell, CtxNrn=cortical neuron, aRG=apical radial glia, oRG=outer radial glia, IP=intermediate progenitor, DL=deep layer, PN=projection neuron, Cfu=corticofugal, CPN=cortical projection neuron.

**NeMO Analytics links in Supplemental Figure legends:**

NeMOlinkS01: <https://nemoanalytics.org/p?p=p&l=NeocortexEvoDevo&c=MammCtxDev.jNMF.p7&algo=nmf>

NeMOlinkS02: <https://nemoanalytics.org/p?l=NeocortexEvoDevo&g=DCX>

NeMOlinkS03: <https://nemoanalytics.org/p?p=p&l=NeocortexEvoDevo&c=MammCtxDev.jNMF.p40&algo=nmf>

NeMOlinkS04: <https://nemoanalytics.org/p?p=p&l=AdultNeoctxLayers&c=HsCtxLayer.jNMF.p20&algo=nmf>

NeMO Analytics Neocortex Development landing page and guide: <https://nemoanalytics.org/landing/neocortex>
