## Supplementary material for "A Curated Compendium of Transcriptomic Data for the Exploration of Neocortical Development": Methods

**Data ingestion, processing and upload to NeMO Analytics**

To expedite analysis, facilitate broad exploration, and maintain reproducible results with original publications, the majority of data in NeMO analytics are formatted as authors originally processed them, most often log transformed and column normalized (e.g. log2(CPM+1) or similar). Original online source repository locations are listed for all individual neocortical datasets in NeMO Analytics [HERE](https://docs.google.com/spreadsheets/d/1xag_73Y1xydcCUnDYwn8rlF2rEbbMdTw/edit?usp=sharing&ouid=104175833608653433199&rtpof=true&sd=true). Processing details and/or code for individual datasets can be provided upon request. Author-curated metadata were utilized to preserve analysis results by original authors wherever available, including cell type labels, lower-rank approximations and de-identified donor/sample metadata. With this highly curated sample metadata (obtained from direct collaboration with original authors when not provided in online repositories), NeMO Analytics enables the custom assembly and simultaneous visual inspection of many low-dimensional maps of datasets from the collection. Harmony or Seurat were used to integrate individual datasets for clustering and/or low-rank approximations/visualizations when needed, e.g. for additional subsetting of large published datasets. Whether derived from a small focused experiment, or the product of “integration” across coordinated studies to create broad molecular atlases, each public data matrix in NeMO Analytics is analyzed and displayed within its own low-dimensional representation, with custom user-created visualizations. This allows assessment of precise signals in large numbers of complex, related datasets without the challenges of forcing diverse experimental data into a single unified dimension reduced space.

We invite researchers to upload their own datasets to explore them in the context of the existing resources in NeMO Analytics and allow others to explore their data as well. We have made tools and tutorials for dataset uploading and gene list uploading, with additional resources for formatting data for NeMO Analytics upload within R at [CarloColantuoni.org](https://www.carlocolantuoni.org/articles/data-collections-at-nemo-analytics).

**Consensus MetaMarker cell type calling**

In order to first establish a coarse consensus cell type labeling across mammalian neocortical development, we used composite expression of MetaMarkers (PMID: 37034757), cell type markers that are robust across many studies spanning cortical regions and developmental time (Figure 1B, colored legend). Usage of MetaMarker gene lists to make consensus cell type calls across datasets is diagrammed in Fig S1. We selected the top 100 MetaMarkers for each cell class using the published marker lists and performed a gene-set-scoring based classification approach to label each cell as a single MetaMarker cell class, allowing for an independent, standardized cell class assignment to overlay on our SJD dimensions. Specifically, for a given dataset, the addModuleScore() function in Seurat (PMID: 25867923) was applied using each MetaMarker cell class gene list. Each cell was assigned to a MetaMarker cell class based on its highest gene set score. In order to focus these analyses on excitatory neurogenesis, cells classified as “Glutamatergic”, “IPC”, “Dividing”, or “NPC” were retained and cells classified as “Nonneural” and “Gabaergic” were removed. Author-labeled cell types that were not part of the excitatory neurogenic trajectory were also omitted from these analyses. This was performed on each of the three mammalian neocortical scRNA-seq datasets in the p7CtxDev and p40CtxDev joint decompositions (Figures 1 and 3).

**Joint Decomposition**

We have developed approaches to joint matrix decomposition in Structured Joint Decomposition (SJD, with implementations in the R programming language and python via rpy2; doi.org/10.1101/2022.11.07.515489, <https://github.com/CHuanSite/SJD>). By defining within-experiment variation that is common to multiple matrices, the SJD framework may avoid technical artifacts resident in individual datasets, as well as technological differences between datasets, while allowing the mapping of common axes of variation without requiring alignment to a single dimension reduced space. SJD can leverage several different matrix decomposition algorithms including, principal component analysis (PCA), independent component analysis, (ICA), and non-negative matrix factorization (NMF) to define shared dynamics across multi-omics matrices. SJD implementations of each of these also enable semi-supervised structured decompositions by incorporating biologist defined groupings of the input matrices that define data subsets expected to share common molecular elements. For example, when decomposing four matrices, two from mouse and two from human, a user can request that SJD define dimensions common to all the matrices, and then also elements specific to each of the two species matrix subsets. Because any matrix decomposition requires a strictly matched row structure (i.e. the same genes/features in each matrix), all decomposition functions in SJD automatically perform orthologue mapping across species using the biomaRt package, so that input matrices can be derived from multiple species.

In this report, we perform three joint decompositions - these are described in detail in the next sections. Resulting gene loadings and related gene enrichment tests for all three joint decompositions are in the Supplemental tables. Code and datasets used as input are at [CarloColantuoni.org](https://www.carlocolantuoni.org/articles/neocortical-development).

**jointNMF decomposition of fetal scRNA-seq data from mouse, macaque, and human neocortex (Figures 1 & 2)**

The same collection of three datasets was used for the two joint decompositions focused on conserved elements of mammalian cortical neurogenesis analyzed in Figures 1 and 3: scRNA-seq data from mid-gestational mouse (PMID: 34321664), macaque (PMID: 37824652) and human (PMID: 34390642) were assembled, incorporating only cells within the excitatory neurogenic lineage. There were a total of 15469 genes with detectable expression and identifiable orthologues across the mouse, macaque and human data matrices. For more details on cell type calling and filtering, see the “Consensus MetaMarker cell type calling” section above. The jointNMF() function in the SJD package in R was applied to these three matrices, requesting 7 dimensions common to all three datasets for the Figure 1 analysis (p7CtxDev), and 40 dimensions for the Figure 3 analysis (p40CtxDev). Non-negative matrix factorization (NMF) avoids the orthogonality constraints required by PCA, and is therefore well-positioned to identify the many distinct, yet highly correlated signals that are common in multi-omic data. While jointNMF() and all the decomposition functions in SJD are capable of doing so, we did not request detection of additional dimensions of variability shared among subsets of the matrices. Using the jointNMF() function without additional structured dimensions is equivalent to concatenating the matrices and running the NMF on this concatenated matrix with additional normalization steps to ensure that each matrix is weighted equally (weighting can also be modified in SJD if desired), accounting for differences between matrices in magnitude of total variance and in sample/cell numbers. The result is an approach that harnesses NMF to identify a single set of dimensions that describe common variation in all the input matrices. A similar approach has been used to identify biologically meaningful elements that are common to expression and DNA methylation data matrices (PMID: 24223768). In addition, the lack of additional structured dimensions in this decomposition allows for maximal interpretability and simpler transfer of the resulting dimensions to other datasets. Notably, we provide links throughout the text where the projection of these dimensions can be inspected across collections in NeMO Analytics (see “Projection analyses” section below). While the projection of more complex structured decompositions are possible using SJD, this requires additional coding and cannot be done online in NeMO Analytics. We invite computational biologists to use SJD to design creative biologically structured *in silico* experiments using diverse biologically linked datasets, along with the accompanying projection methods in SJD to take advantage of these possibilities (<https://github.com/CHuanSite/SJD>).

**jointNMF decomposition of human adult neocortex snRNA-seq data (Figures 4 & 5)**

For the neuronal decomposition explored in Figures 4 & 5, only excitatory neurons from layer-specific microdissected adult human neocortex snRNA-seq data generated using the SMART-seq V4 technology were used (based on original author cell type calls). Again here we applied the jointNMF() approach, requesting 20 dimensions (p20CtxLayers), to define shared molecular dynamics across 5 matrices (each containing data from 1 donor), spanning a total of 8 neocortical regions in 2 studies: Jorstad 2023 (PMID: 37824655; 3 donors) and Bakken 2021 (PMID: 34616062; 2 donors) - see Figure S4 for more details and full visualization of the resulting transcriptomic signatures. There were a total of 14272 genes with detectable expression in all 5 snRNA-seq data matrices from adult human neocortex. Here, again we requested only components shared across all matrices, with no additional structured dimensions in matrix subsets to facilitate projection and interpretation.

**Gene set enrichment**

For each of the three decompositions performed (see “Joint Decomposition” section above), we performed tests of enrichment on 4 different collections of gene sets 1] MAGMA gene-level analysis of GWAS results from neuropsychiatric disorder and brain structure studies (PMID: 25885710; 9 GWASs tested for significance) , 2] Custom disease gene lists of high penetrance, low frequency variants for specific cortical disorders (PMID: 38915580; 106 disease gene lists tested, 3] Kyoto Encyclopedia of genes and genomes (KEGG; https://www.genome.jp/kegg/; 186 gene lists tested), and 4] Gene Ontology (GO; https://geneontology.org/; 10,532 gene lists tested). The MAGMA analyses are described is the “MAGMA enrichment analyses” section below. To perform enrichment analysis of disease, KEGG and GO gene lists in each of the three jointNMF gene loading matrices, we employed the wilcoxGST() function from the limma package in R. Because gene loadings from NMF are all positive, we used an F-distribution null with a mixed alternative hypothesis. All p-values are uncorrected. Results from all tests for each of the joint decompositions performed in this report are in Supplemental Tables 1-3. Selected results are listed in Figures 1 & 3. Code at [CarloColantuoni.org](https://www.carlocolantuoni.org/articles/neocortical-development).

**MAGMA enrichment analyses**

Gene loadings from each jointNMF decomposition were tested for enrichment of gene-level risk estimates from GWAS summary statistics calculated using MAGMA (PMID: 25885710) from recent studies on neuropsychiatric disease (PMID: 35396580, PMID: 34002096, PMID: 30804558), and brain MRI structural phenotypes in the UK BioBank. Summary statistics for SCZ, BD, and ASD GWAS were downloaded from the Psychiatric Genomics Consortium (PGC) portal (<https://pgc.unc.edu/for-researchers/download-results/>; PMID: 35396580, PMID: 30804558, PMID: 34002096). UK Biobank MR brain GWAS summary statistics were downloaded from the Oxford Brain Imaging Genetics Server (<https://open.win.ox.ac.uk/ukbiobank/big40/>; PMID: 33875891). MAGMA gene-level analysis using gene covariate mode with decomposition loadings as covariates was run on the summary statistics using default parameters (PMID: 25885710). All MAGMA analyses were performed using NCBI37.3 gene locations.

**Projection analyses (transfer learning)**

In this report we have laid out a framework that first uses joint decomposition of related multi-omic matrices to define shared molecular change and then employs transfer learning methods, i.e. using projection or weighting matrices trained on other data, to more broadly explore the functional biology of these latent spaces across additional biologically linked data. The schematic below is a map of the datasets, decompositions and projections which we use throughout this report.

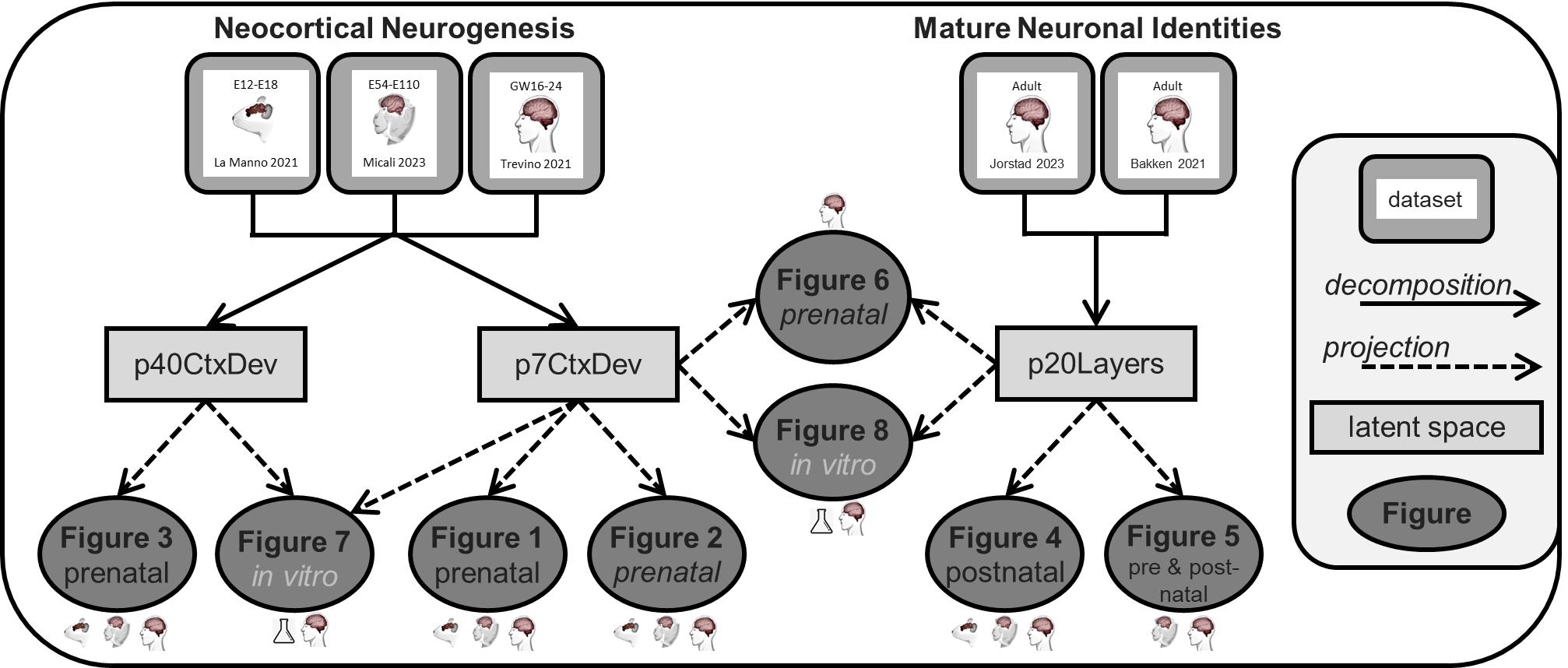

Examining a dimension of variation defined by NMF within a new dataset is a form of projection onto the linear space spanned by the NMF columns, which are not typically orthogonal principal components, however, there are multiple approaches suitable to estimate embeddings for single cells in the new dataset. All of the projections visualized in this report were made with least-squares optimization to estimate sample embeddings in new data given gene loadings from the jointNMF decompositions. This method is implemented in the projectR package in R (PMID: 32167521; <https://www.bioconductor.org/packages/release/bioc/html/projectR.html>). A second method for estimating new sample embeddings from an NMF gene loading matrix by running the NMF algorithm with a fixed loading matrix has been implemented in the SJD package (<https://doi.org/10.1101/2022.11.07.515489>, <https://github.com/CHuanSite/SJD>). Both of these methods are available online in NeMO to estimate and visualize NMF projections. All of the NMF projections links in this report point to projections using least-squares optimization. However, simply replacing the “algo=nmf” with “algo=fixednmf” in any projection link will switch the projection method. Methodology for the projection of PCA is also implemented in NeMO Analytics (using “algo=pca”). In addition to projecting data using PCA gene loadings, this can serve as a multi-purpose projection method as it computes the sum of expression of (gene expression levels) * (gene loadings) for each cell/sample. Finally, also in NeMO, there is a transfer method for simple, unweighted gene lists that is well suited to single-cell RNA-seq analysis. This method simply counts the number of non-zero expression levels in the list of genes given. Code projection analyses can be found at [CarloColantuoni.org](https://www.carlocolantuoni.org/articles/neocortical-development). We invite researchers with specific gene signatures of known interest to upload their gene lists or gene loadings for transfer learning experimentation in the NeMO Analytics environment (tools and tutorials for gene list upload and more at [CarloColantuoni.org](https://www.carlocolantuoni.org/articles/data-collections-at-nemo-analytics)).

Projection analyses are powerful tools to explore transcriptome dynamics discovered in a reference dataset within new target datasets of interest that may be biologically related, but evolutionarily or technologically different (PMID: 32167521, doi.org/10.1101/2022.11.07.515489, PMID: 31121116, https://doi.org/10.1101/2021.03.17.435870). It must be noted, however, that this approach has some very specific limitations. Due to differences in the nature of samples and their preparation and in measurement technologies (the same issues that make integration methods impractical across heterogeneous datasets), effects observed in transfer learning are relative within each projected dataset, i.e. units resulting from projection analyses are comparable only within, not across, projected experiments. For this reason, in this report we display all data projections on a minimum to maximum scale bounded by each individual projected dataset.

This quality of transfer learning analysis across disparate datasets brings up the important point that there will always be a maximum and a minimum in a projection of any dataset, and we must take care in interpreting the relative scale of these signals, without formal statistical inference. While these methods can be used to approximate cell types across studies with similar cell type distributions, they are not cell deconvolution or cell type calling per se (such as <https://portal.brain-map.org/atlases-and-data/bkp/mapmycells> or <https://azimuth.hubmapconsortium.org/>). Our approach is more general, measuring the strength of a transcriptomic signal defined in one dataset within a new dataset of interest. For example, the cells in dataset 2 with the highest levels of a transcriptomic signature defined in cell type A in dataset 1, are NOT necessarily cell type A. They are simply the cells in the new experiment with the highest levels of the cell type A signature. While this is a limitation of projection analysis, it is also a strength of these methods if biological care is taken in the interpretation of projected signals. This can enable the discovery of relationships across diverse cell types and states, e.g. cell type B express may express high levels of a cell type A expression signature at a particular point in development.

Several recent reports use similar decomposition or transfer learning strategies to explore new data, although most are more tailored to specific tasks (e.g. mapping new scRNA-seq data to a reference, or removing batch effects), while we aim to leverage these methods to gain broad perspectives on heterogeneous datasets. These are many and include: ExpiMap (PMID: 36732632), scArches (PMID: 34462589), treeArches (PMID: 37502708), scGAD (PMID: 36869836), BEENE (PMID: 37561107), Symphony (PMID: 34620862), scMC (PMID: 33397454), SIMS (PMID: 38823397), scCross (PMID: 39075536), scBOL (PMID: 38678389), MIDAS (PMID: 38263515), scPML (PMID: 38097699), scMRA (PMID: 34623390), CIForm (PMID: 37200157), ABC (PMID: 38213820), MOFA (PMID: 32393329 and PMID: 29925568), transfer learning for scRNA-seq clustering (PMID: 31889137), and a new review of reference mapping (PMID: 38729109). Similarly, our approach is distinct from integration methods which seek to map samples or cells from multiple datasets into a single dimension reduced space, such as Seurat (PMID: 34062119 and PMID: 31178118), Harmony (PMID: 31740819), scDART (PMID: 35761403), scAWMV (PMID: 36383176), and LIGER (PMID: 31178122). Of these many approaches, perhaps scArches (PMID: 34462589) and MOFA (PMID: 32393329 and PMID: 29925568) are most closely related to our techniques here. In contrast, our decomposition and projection strategy leaves samples from each dataset in their own space, but arranged along a shared axis of gene expression variation. Importantly, the true origin / zero point along this axis is not the same across datasets, so, as stated above, units of projected embeddings are comparable only within datasets and we gain insight by comparing relationships between samples in a study to relationships between samples in another.

**Trevino et al 2021 Cell type reannotation and CellOracle analysis**

This analysis of the Trevino 2021 data is very similar to the analysis conducted in Mato-Blanco 2024 (PMID: 38915580). Specifically for the CellOracle analysis in Figure 3, we re-annotated cell types in the Trevino 2021 data: Single-cell RNA and ATAC data of prenatal human cortical development (PMID: 34390642) were obtained from the Gene Expression Omnibus (GEO) dataset GSE162170. We used scanpy (PMID: 29409532) to preprocess the expression data. We retained cells with less than 10% mitochondrial counts and genes with at least one count. This resulted in a dataset of 53,231 cells and 27,886 expressed genes. To classify cell-cycle phases in each cell, we scored the expression of cell-cycle phase-associated genes (PMID: 27124452) and classified cells based on these genes. Next, we identified highly variable genes within each donor and selected the top 5,000 genes. We normalized the data to a value of 1e4 counts per cell, log-transformed it, and scaled it. To model gene expression and integrate data across samples, we employed scVI (PMID: 30504886), a deep generative model for single-cell RNA-seq data analysis. We used sequencing batch as a batch variable and included mitochondrial and ribosomal count fractions as covariates, as well as cell-cycle scores. The latent space size was set to 10, and we obtained corresponding embeddings for all cells in the preprocessed data. To further define early cell types present in the dataset, we selected progenitors (cycling, multipotent glial (mGPCs), oligodendrocyte intermediate (OPCs) and neuronal intermediate (nIPCs) and radial glia (early, late, and truncated) cells. We used 15 nearest neighbors in the embedding space, created a UMAP representation (https://arxiv.org/abs/1802.03426), and identified cell clusters using the Leiden algorithm (PMID: 30914743). Using a selection of known marker genes (PMID: 34390642, PMID: 26406371), we assigned cell type identities based on the clusters’ expression and excluded a low-count cluster, ependymal cells and interneurons. In this way, we could identify ventricular, truncated and outer RGCs, as well as early neurons, nIPCs, mGPCs, OPCs and astrocytes.

For the CellOracle analysis, using the reannotated data, we selected RGCs (ventricular, truncated and outer), as well as mGPCs, astrocytes and OPCs to obtain a representative dataset of the gliogenic differentiation and maturation of RGCs. We preprocessed the data as suggested in the CellOracle pipeline (PMID: 36755098). In brief, genes with 0 counts were removed, count data was normalized per cell and only the 3000 top highly variable genes were used. Additionally, we forced the inclusion of a selection of genes of interest, i.e., HOPX, FOXN3, FOXJ2, NAB2, HES2, ATXN7L3, FOXD2, MAFG, ZNF423, PCGF1, NR1H4, MIF1, UBN1, TFEB, NR0B1, ICSBP, IRF8 and MIF. Then data were normalized, log-transformed and scaled again. We computed k-nearest neighbors (knn) with 10 neighbors in scVI embedding space and created a diffusion map of 20 components using these neighbors.knn with 20 neighbors was computed again on the diffusion map and a ForceAtlas2 (PMID: 24914678) representation was created. For CellOracle's pseudotime computation, we selected knn as the method to fit with 3 neighbors and we manually selected a cell from the earliest sample and low pseudotime. Sample-matched scATAC-Seq data (PMID: 34390642) were leveraged to reconstruct a base gene regulatory network (GRN). Monocle3 (PMID: 30787437) was used to preprocess the data using LSI as the normalization method. Peaks and peak-to-peak co-accessibility were obtained by Cicero (PMID: 30078726). Peaks overlapping a transcription-start site (TSS) were annotated to the corresponding gene and only those peaks with a co-accessibility greater than 0.8 were retained. Peaks were scanned for motifs using CellOracle’s scan function, which uses the gimmemotifs (doi.org/10.1101/474403) motif scanner (fpr: 0.02, default motif database: gimme.vertebrate.v5.0 using binding and inferred motifs, and cumulative binding score cutoff: 8) to generate an annotated peak-motif binary matrix which is the base GRN in CellOracle. The computed base GRN was fit to each cell type using the CellOracle functions: first, cell type specific links were retrieved by fitting the GRN to the cell-type specific expression matrix using a bagging ridge regression model (bagging_number: 20, alpha: 10). The links in the resulting networks were filtered by p-value (< 0.001) and from those, only the top 2000 links were retained based on their mean coefficients. After this filtering, the model is fit once more to adjust the coefficients of the preserved links. Before CellOracle's simulation can be performed, a simulation grid needs to be fit in the cell pseudotime map to estimate a developmental flow in the data. For this, we manually selected the following parameters in the CellOracle pipeline: ridge alpha: 10, mass smooth factor: 0.8, grid points: 40, minimum mass: 0.0017, fit method: knn with 200 neighbors, scale of the flow: 40. These parameters were then used to perform the simulation step as well, combined with the desired gene and expression value to simulate. We analyzed the effect of completely knocking-out the expression of a gene, i.e., simulating an expression value of 0, and the effect of overexpressing it in the whole data, i.e., simulating a constant expression value of double the observed maximum. We simulated the knock-out and overexpression of all transcription factors available in the expression data and GRN. The results represent the cell-type transitions observed in CellOracle's simulation (run for 500 steps, replicated 5 times).
